# Scaling and foraging behavior drive the evolution of humeral shape in hummingbirds

**DOI:** 10.1101/2025.08.11.669688

**Authors:** Juan Camilo Ríos-Orjuela, Carlos Daniel Cadena, Alejandro Rico-Guevara, Ilias Berberi, Lauren Miner, Roslyn Dakin

## Abstract

Understanding how locomotion-related skeletal elements evolve under biomechanical and ecological constraints is central to animal evolutionary biology. In hummingbirds (Trochilidae), the humerus plays a key role in force transmission during hovering and flapping flight, yet the drivers of its shape evolution have not been examined. We combined geometric morphometrics with phylogenetic comparative analyses to examine humeral shape variation, evolutionary rates, and phenotypic integration in male hummingbirds from 78 species. Our analyses identified humerus allometry as the dominant predictor of shape, revealing a pattern in which larger humeri show broader proximal epiphyses, increased shaft robustness, and reduced curvature. In addition to this scaling pattern, we find that male humerus shape differs subtly among hummingbird species that differ in the use of aggression during nectar foraging. Evolutionary rates of humeral shape were heterogeneous and decoupled from ecological predictors. We also find phenotypic integration between the proximal and distal regions of the humerus, indicating coordinated evolution. Together, these results show that humeral evolution in hummingbirds is governed primarily by biomechanical scaling and internal integration, with foraging ecology introducing secondary, size-dependent modifications. This work highlights the importance of considering scaling and internal integration when interpreting morphological evolution in locomotor systems across vertebrates.

## Introduction

Among skeletal elements, the avian humerus plays a central role in flight mechanics and ecological adaptation, connecting the axial skeleton to the forelimb and anchoring the primary muscles driving the wingbeat [1]. Variation in humeral morphology is expected to influence wingbeat kinematics and, indirectly, the magnitude and direction of aerodynamic forces, as well as flight control and maneuverability [2]. Despite this biomechanical importance, comparative studies of avian flight have largely focused on external features such as wing aspect ratio or wing loading [3], which capture overall aerodynamic performance, but fail to reflect underlying skeletal variation and musculoskeletal function.

In hummingbirds (family Trochilidae), the humerus is short and robust, reflecting the extreme mechanical demands of their specialized flight mode, characterized by sustained hovering, high wingbeat frequencies owing to their small body size, and rapid wing rotation throughout the stroke [2]. Unlike the long, slender humeri typical of birds adapted for gliding or forward-powered flight [4], the hummingbird humerus functions as a rigid lever transmitting large forces from enlarged flight muscles that drive lift production and rapid wing rotation [5]. This configuration enables precise control during hovering and agile maneuvers but also imposes high mechanical loads [2]. As a result, compared to other birds, the hummingbird humerus exhibits increased cortical thickness, a relatively straight shaft, and pronounced muscle attachment sites at the deltopectoral crest (e.g., Fig. 1A; [6]). This morphology reflects a biomechanical trade-off between minimizing inertial resistance during rapid wing movements and maintaining the structural stiffness required to transmit large aerodynamic forces. Such opposing demands are expected to strongly constrain humeral shape, making it a key element for understanding the evolution of flight-related morphology in hummingbirds. Despite the functional centrality of the humerus, and the exceptional diversity of hummingbird family (>350 species), comparative analyses of its evolution across this group have not been examined.

**Figure 1.**
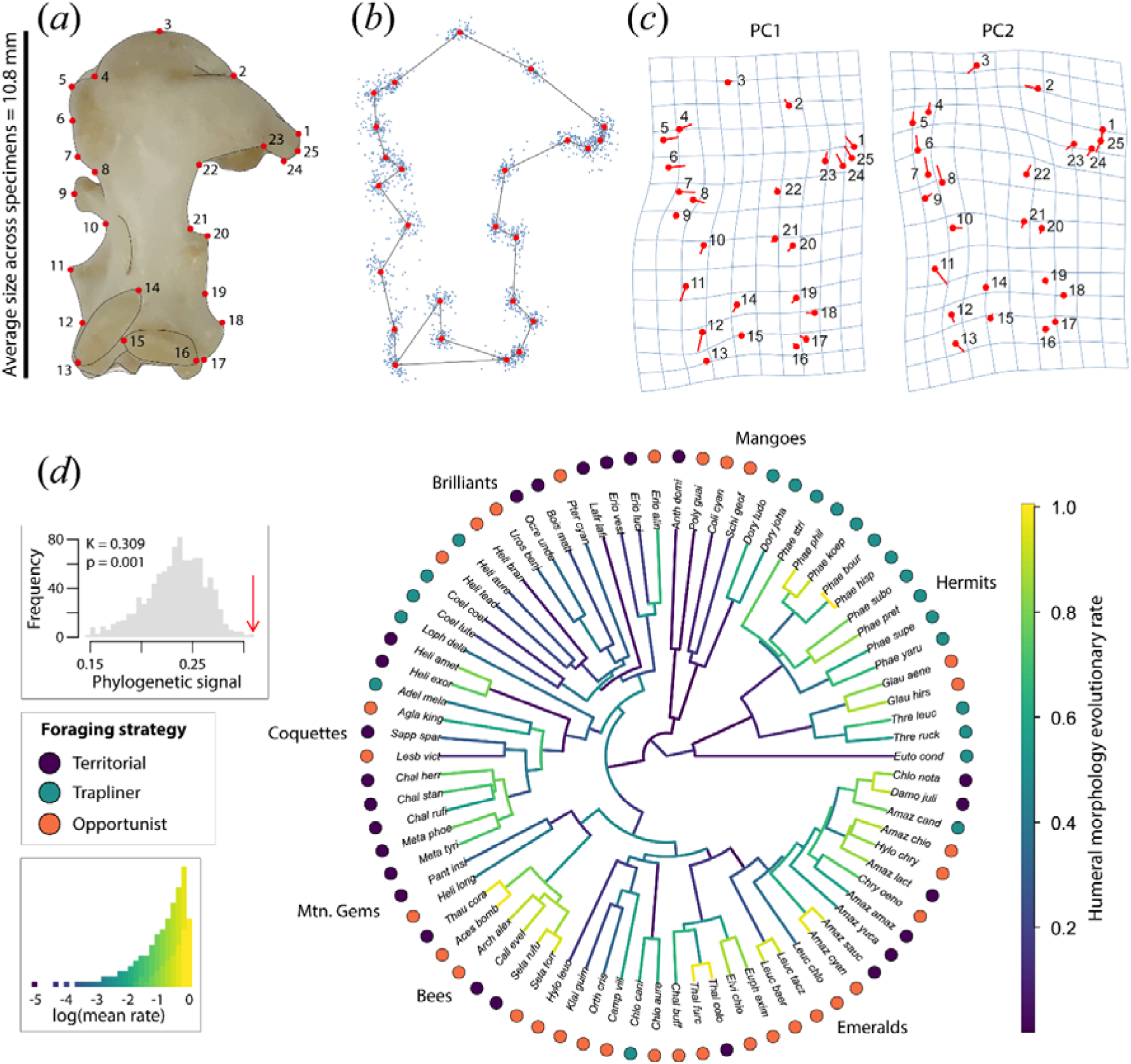
Hummingbird humeral shape shows strong phylogenetic structure and evolutionary rate heterogeneity across major clades. (a) Right humerus illustrating the 25 anatomical landmarks used to quantify humerus shape, averaged across male specimens and scaled to mean size. (b) Procrustes-aligned landmark dispersion across specimens, illustrating overall humeral shape variation within the hummingbird family. (c) Thin-plate spline deformation grids for PC1 and PC2, showing the major axes of humerus shape variation captured by our morphometric analysis. (d) Phylogenetic distribution of humerus shape evolutionary rates across the 78 hummingbird species analyzed; branch colors indicate estimated evolutionary rates, and tip symbols denote foraging strategy. Insets show the null distribution of phylogenetic signal with the observed value indicated by an arrow, and the distribution of log-transformed mean evolutionary rates.

From a biomechanical perspective, the evolution of load-bearing skeletal elements is constrained by scaling relationships between size, force production, and structural resistance. Under geometric similarity, body mass scales with linear dimensions cubed, whereas resistance to bending or axial loads scales with cross-sectional area and thus with linear dimensions squared [7]. Maintaining constant mechanical stress therefore requires load-bearing bones to scale at greater than isometry. This expectation applies broadly to locomotor elements, including the humerus, which experiences substantial bending and torsional loads during the wingbeat. Even in the absence of ecological differentiation, fundamental biomechanical principles predict a disproportionate increase in humeral robustness with body size.

The above general biomechanical expectations are further modified by the exceptional allometric scaling of the wing in hummingbirds, a group where species vary ∼10-fold in body mass. Unlike most birds, larger hummingbird species exhibit disproportionately large wings relative to body mass, and unusually low wing loading [8–10], a departure from isometry that is central to sustained hovering flight. As a result, aerodynamic and inertial forces acting on the wing do not increase proportionally with body mass alone. Because these forces are transmitted proximally through the wing skeleton, the humerus must accommodate both increases in body size and the disproportionate scaling of wing dimensions. Consequently, humeral morphology reflects an interaction between general biomechanical constraints and clade-specific wing allometry, making hummingbirds an informative system for studying size-related skeletal evolution.

Hummingbirds also exhibit species differences in foraging behavior that are linked to the use of different flight modes. In some species, males primarily defend floral territories, engaging in aggressive chases, rapid accelerations, and tight turns, whereas other species adopt traplining strategies involving predictable routes and sustained flight among dispersed resources [9,11]. These differences in the use of aggressive behavior while foraging are expected to impose different biomechanical demands on the flight apparatus. Territorial hummingbirds may benefit from shorter, more robust humeri that enhance force production and maneuverability [2], whereas traplining behavior may favor longer, more slender humeri associated with sustained flight efficiency [12,13]. However, foraging strategies are coarse ecological proxies and may be confounded with body size, habitat structure, and sex-specific behaviors. In many hummingbird species, males and females differ in morphology and flight performance, often as a result of sexual selection for enhanced agility in males [14–17]. Here, we focus primarily on humerus evolution in males, but we present measurements from a smaller sample of females in the Supplementary Material.

Beyond foraging strategy, additional species differences related to the flight environment and larger-scale movements may also shape humeral morphology. Habitat density modulates maneuvering demands, with closed environments requiring frequent turning and fine-scale control, whereas open habitats are expected to favor fast flight and straighter flight paths [18,19]. Migratory behavior represents another axis influencing morphological evolution, from resident species to those undertaking seasonal migrations that require sustained long-distance flight [20]. Broad environmental gradients such as elevation, latitude, and geographic range size may further shape humeral morphology. High-elevation environments impose hypoxia and reduced air density, increasing the mechanical and metabolic demands of flight and driving divergence in wing shape, body size, and metabolic traits in hummingbirds [21–23]. Latitude and geographic range size similarly capture variation in climatic heterogeneity, which may influence dispersal and exposure to diverse flight conditions [24,25].

Given this background, we expect humeral morphology in hummingbirds to reflect a complex interplay among biomechanical scaling, ecological demands, and evolutionary history. However, the extent to which ecological variables predict humeral shape independently of body size, and whether they are associated with differences in evolutionary rates, remains underexplored [26,27]. In this context, evaluating evolutionary rates provides a complementary axis for testing ecological effects beyond static shape differences, as accelerated shape evolution may reflect ecological opportunity or functional innovation, whereas low rates may indicate stabilizing selection or strong biomechanical constraints [28,29].

The hummingbird humerus, with distinct proximal and distal functions, also provides an opportunity to investigate phenotypic integration. Proximally, the humerus anchors the pectoralis and supracoracoideus muscles that power the downstroke and upstroke, whereas distally it articulates with the radius and ulna, transmitting forces that shape wing motion and orientation [1,30]. Understanding whether these regions evolve in a coordinated manner offers insights into developmental constraints and adaptive flexibility [31–33]. Locomotor traits rarely evolve independently [34], instead exhibiting integration or modularity driven by shared developmental pathways and functional covariation [35,36]. While most studies examine integration across multiple skeletal elements, analyzing integration within a single bone can reveal how functionally distinct regions remain evolutionarily coordinated despite localized biomechanical specialization.

Here, we investigate how biomechanical scaling, species ecology, environmental gradients, and evolutionary processes jointly shape humeral morphology in male hummingbirds. First, we tested whether humeral shape exhibits strong allometric scaling with body size. In this clade, wing length increases more than expected from isometric scaling with body mass (approximately mass^0.5), resulting in disproportionately long wings in larger species [8–10]. Because the humerus transmits aerodynamic and inertial forces from the wing to the body, we predict that humeral robustness exhibits positive allometry (i.e., scales greater than isometry) with body size. We further assessed whether allometric variation in humeral shape is associated with differences in evolutionary rates, recognizing that size-related constraints may promote directional change or impose stabilizing selection.

Second, we examined whether ecological context modulates humeral morphology and evolutionary rates beyond size and allometry. We hypothesize that differences in foraging behavior will impose distinct biomechanical demands: we expect territorial species to show shorter, more robust humeri consistent with enhanced force transmission during high-power flight, whereas traplining species may exhibit more elongated humeri associated with sustained flight efficiency [9,37]. We further predicted complementary effects of habitat structure and migratory strategy, with closed habitats favoring greater maneuverability and open habitats favoring straighter flight trajectories; we expect migratory species to exhibit morphologies linked to sustained directional flight [12,18,38]. In contrast, when examining evolutionary rates, we did not assume the ecological predictors would have a single directional effect, as selection may promote innovation or impose stabilizing constraints depending on ecological variability and biomechanical trade-offs [39].

Third, we assessed whether broad environmental gradients shape humeral morphology and evolutionary rates. High elevation imposes hypoxia and reduced air density, increasing aerodynamic and metabolic demands. We therefore predicted that elevation may be associated with shifts in humeral morphology, whereas its effects on evolutionary rates could arise either through increased selective pressure promoting faster change, or through environmental constraints limiting phenotypic divergence. Latitude and species’ geographic range size are broad descriptors of environmental variation, and we accounted for these in our analyses, although we did not have a priori directional expectations for these two predictors.

Lastly, we hypothesized strong internal phenotypic integration in the hummingbird humerus, with coordinated evolution between proximal and distal regions due to their shared role in force transmission and wing mechanics. This is expected because modifications in one region (e.g., altering muscle attachment sites) would require compensatory changes to maintain flight performance. However, bones may also evolve modularly when regions face distinct functional demands [34,40,41]. Whether such modularity also occurs within the hummingbird humerus remains an open question, with implications for how evolutionary change may be distributed across the structure. By integrating ecological predictors and morphometric analyses within a phylogenetic framework, we provide a comprehensive assessment of the drivers of humeral morphological evolution.

## Methods

### Specimens and image acquisition

We imaged humeral morphology of skeletal specimens from 80 hummingbird species (∼22% of extant Trochilidae with 7 of 9 clades represented in our sample [42]) at the Louisiana State University Museum of Natural Science (LSUMNS, see Table S1 for specimen details). Sampling was uneven due to limited skeletal material: ∼87% of species (70/80) were represented by a single male specimen. We focused our main analyses on male specimens to minimize variation associated with sexual dimorphism and to ensure sampling consistency, as male skeletal material is more widely available [43]. We provide a preliminary characterization of a smaller sample of females in the supplement.

The right humerus was photographed in cranial view with a scale bar (0.1 mm divisions) using a Canon EOS M50 (6000 × 4000 px) with a 28 mm macro lens. Images were acquired with orthogonal orientation and fixed camera distance using a copy stand to ensure cross-species comparability [44].

We digitized 25 homologous anatomical landmarks per humerus, primarily along the outline (Fig. 1A–B), using TPSDig2 v2.32 [45]. Landmarks were selected for anatomical consistency and biomechanical relevance; semilandmarks were not used (see Table S2 for details). We screened each photograph for humerus orientation, planar alignment, and landmark visibility and excluded specimens that failed to meet these criteria. Two species (*Calypte anna* and *Goethalsia bella*) were excluded because the specimens (one individual each) did not meet landmark quality criteria. After screening, 78 species remained for downstream analyses.

To assess the potential influence of geometric uncertainty on the morphometric dataset, we considered several non-statistical sources of measurement error. Image resolution introduces ≤1–2 pixels of landmark uncertainty (∼0.1% of humeral length), intra-observer digitization error is typically 0.5–1.5%, inter-observer error is negligible (single observer), and local anatomical ambiguity may add ∼1–3% variability. Combined uncertainty (∼2–4% of inter-landmark distances) is well below interspecific shape differences and unlikely to affect major axes of variation or phylogenetic patterns.

### Morphometric data preparation

We conducted all analyses in R 4.4.2 [46]. We aligned landmark configurations using Generalized Procrustes Analysis (GPA) to remove effects of position, orientation, and scale [47], using gpagen() in *geomorph* [48]. For species represented by more than one specimen, we used the species-average humerus shape in all subsequent analyses. We used principal component analysis (PCA) of the Procrustes-aligned coordinates to visualize overall shape variation and morphospace structure (Fig. 1C). We did not use PCA scores for hypothesis testing.

### Species traits

For each species, we compiled ecological, geographic, and morphological predictors. Species body mass (g) from Tobias et al., (2022) [49] was included as a proxy for biomechanical scaling, given its relationship with aerodynamic force production. We compiled estimates of species geographic range size (km²) and range centroid latitude (absolute value) from Tobias et al., (2022) [49], and species maximum elevation (m) from Quintero and Jetz (2018) [50]. Body mass and range size values were log-transformed prior to analysis to improve linearity and reduce heteroscedasticity. As a measure of humerus size, we used landmark centroid size, a matrix-derived measure based on Procrustes analysis [51].

Each species’ primary foraging strategy was classified into three categories—territorial, trapliner, and opportunist—using Feinsinger’s framework [52] and Rombaut et al., (2022) [53]. According to this framework, we classify species as territorial if the males primarily defend spatially stable floral patches and exclude competitors through aggressive interactions, whereas species are classified as trapliners if the males primarily visit dispersed flowers and generally do not defend feeding territories. We classify species as opportunists if the males that exhibit flexible or context-dependent foraging behavior (i.e., species that are described as using territorial or non-territorial foraging, depending on the location or conditions). For each species, we manually reviewed foraging strategies assigned in Rombaut et al., (2022) [53] and the corresponding species account in *Birds of the World* [54] to ensure consistency, and to classify species that were not included in Rombaut et al., (2022) [53] (see Table S3 for details).

Habitat density was obtained from Tobias et al., (2022) [49], with each species categorized as using dense, semi-open, or open habitats. Species migratory behavior was classified into four categories: (i) resident, (ii) altitudinal migrant, (iii) short-distance latitudinal migrant, and (iv) long-distance latitudinal migrant, based on species accounts in *Birds of the World* [54], BirdLife International [55], and Barçante (2017) [56]. We considered a species to be migratory (ii - iv) only when explicit evidence of documented movement was available.

### Phylogenetic signal

We assessed phylogenetic signal in humeral shape using Blomberg’s multivariate K statistic [33,57], using physignal() in *geomorph*, based on 1,000 permutations. All analyses used an ultrametric phylogeny from McGuire et al., (2014) [58] pruned to the final species set.

### Phylogenetic modeling of humeral shape variation

To evaluate how ecological and morphological predictors influence humeral shape, we used a phylogenetic generalized least squares (PGLS) framework using procD.pgls() in *geomorph*, with Pagel’s λ estimated by maximum likelihood in the final model [59].

We first evaluated a set of *a priori* candidate ecological models (Table S4) and then fitted a final multivariate model including humerus centroid size, body mass, foraging strategy, migratory status, habitat density, geographic range size, range centroid latitude (absolute value), and maximum elevation. Categorical predictors were coded using sum-to-zero contrasts so that tests reflect overall deviations of each factor from the grand mean rather than comparisons to a single reference level. We verified that collinearity among predictors was low (Pearson’s |r| < 0.36; all adjusted GVIF < 1.5, Table S5). Statistical significance was assessed using 1,000 permutations of residuals under a phylogenetic correlation structure. Because sequential (Type I) sums of squares are sensitive to predictor order (Table S6), we based our inferences on Type III sums of squares, providing order-invariant tests of each predictor’s partial effect while controlling for all others.

We evaluated the sensitivity of PGLS analyses to assumptions about phylogenetic signal (Pagel’s λ) by refitting the final additive model across a range of fixed λ values (0–1) (see supplement Table S7 for details). We performed additional sensitivity analyses to consider interactions between species size and foraging behavior using the same inferential framework (Table S8).

### Discriminant analysis of humeral shape by foraging strategy

To test whether humeral shape differs among foraging strategies, we conducted a Canonical Variates Analysis (CVA) using size-corrected Procrustes coordinates. Allometric effects were removed by regressing the aligned shape coordinates on log centroid size and using the residuals as inputs [51]. CVA was implemented with CVA() in *Morpho* [60], with foraging strategy as the grouping factor, and group separation was evaluated using 1,000 permutation tests. Phylogenetic structure in canonical space was visualized by projecting species scores onto the phylogeny using *phytools* [61].

### Evolutionary rates of humeral shape

We tested whether humeral shape has evolved at different rates in hummingbirds with different ecological traits using compare.evol.rates() in geomorph with 1,000 permutations. We conducted these rate comparisons for the following categorical predictors: foraging strategy, habitat density, and migratory behavior, by evaluating whether group-specific mean rates deviated from expectations given the phylogeny and the distribution of shape change estimates across species.

For continuous predictors, we tested associations between evolutionary rates and each predictor using search.trend()in *RRphylo* [62], which fits phylogenetically informed regressions between branch-specific rate estimates and predictor values. We applied standard *RRphylo* diagnostics to identify potential outliers using the built-in functions *RRphylo*::outliers() and *RRphylo*::search.trend(), summarized in Table S9.

We additionally modeled species-specific evolutionary rates as a univariate response using phylogenetic generalized least squares (PGLS) fitted with *nlme*::gls() under a Brownian-motion correlation structure derived from the same ultrametric phylogeny. Each predictor was evaluated in a separate univariate model.

### Phenotypic modularity and integration

To evaluate phenotypic integration within the humerus, we partitioned the Procrustes-aligned landmark data into two functional modules defined based on avian flight anatomy; we refer to this as the *a priori* modularity hypothesis. According to this hypothesis, the proximal module includes the humeral head, tubercles, and associated articular surfaces involved in shoulder rotation and load transfer during both hovering and flapping. The distal–shaft module comprises the diaphysis, trochlea, capitulum, and distal muscle insertion sites, which primarily mediate force transmission, wingbeat mechanics, and resistance to bending loads during forward flight [30,63]. This subdivision reflects the well-established biomechanical differentiation between the articulation region of the humerus and its load-bearing shaft. We quantified evolutionary covariation between the *a priori* modules using phylogenetic partial least squares implemented with phylo.integration() in *geomorph*, assessing significance of modularity with 1,000 phylogenetically informed permutations. We did not perform alternative modularity searches (e.g., EMMLi-based optimization) because our objective was to test an anatomically motivated prediction of integration rather than explore the full modular landscape of the humerus.

## Results

### Phylogenetic signal

Humeral shape exhibited a significant phylogenetic signal (K = 0.309, p = 0.001, Fig. 1D), indicating that closely related hummingbird species tend to share similar humeral morphologies. The geometric configuration used to quantify humeral shape revealed consistent variation in landmark positions across species, highlighting major axes of morphological differentiation (Fig. 1A–C).

### Predictors of humeral shape variation

The multivariate PGLS analysis identified humerus centroid size as the strongest predictor of humeral shape variation (R² = 0.098, Z = 3.26, p = 0.001; Table 1; Fig. 2). Foraging strategy also showed a significant association with humeral shape (R² = 0.034, Z = 1.98, p = 0.025). Several additional predictors exhibited marginal effects: body mass (R² = 0.017, p = 0.070), migratory behavior (R² = 0.045, p = 0.058), maximum elevation (R² = 0.015, p = 0.095), and absolute latitude (R² = 0.014, p = 0.112) each explained small proportions of shape variation, but did not reach statistical significance. We did not detect effects of habitat density (R² = 0.018, p = 0.394), or geographic range size (R² = 0.010, p = 0.369) on humeral shape.

**Figure 2.**
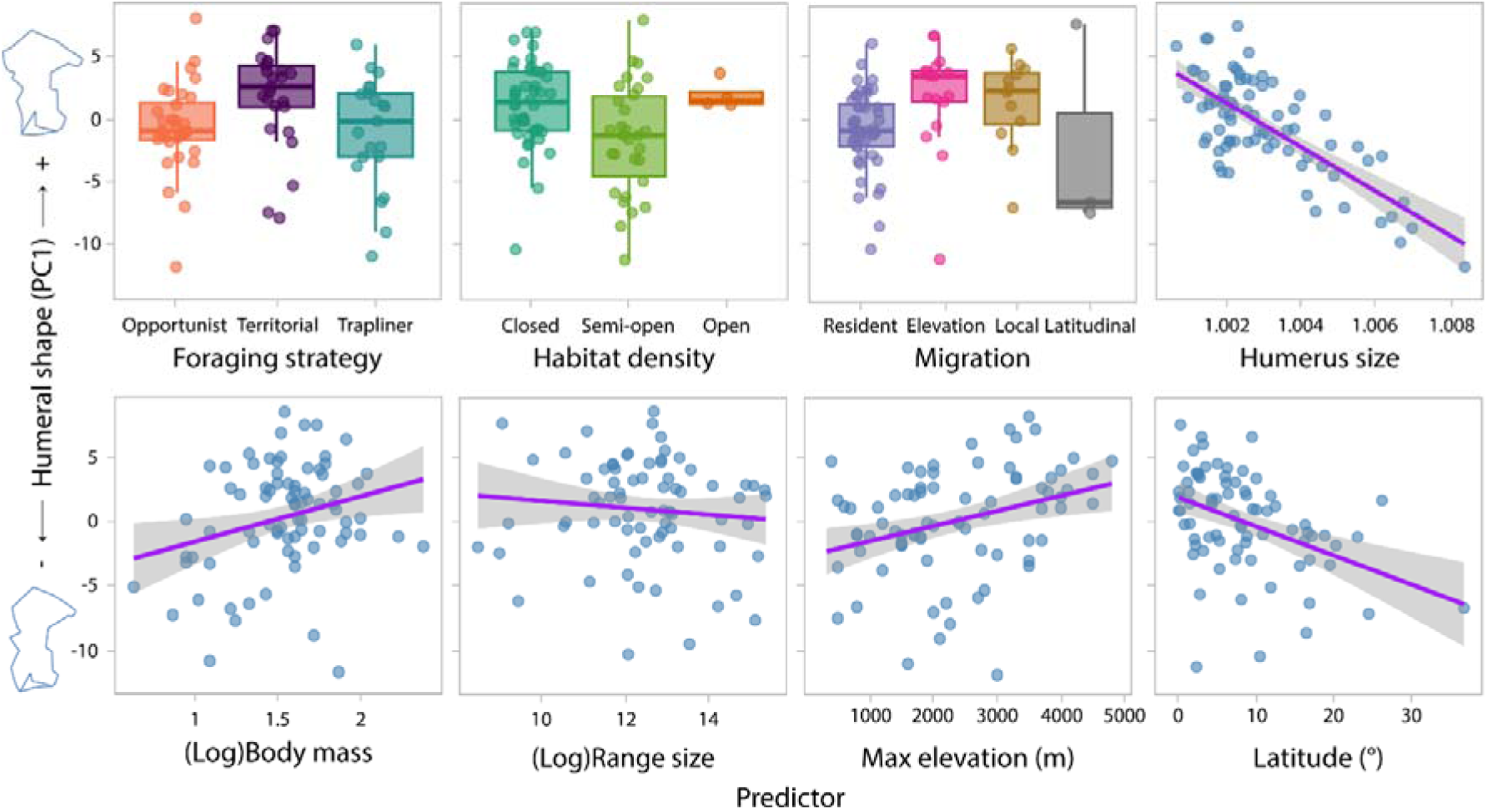
Humerus shape variation depends on humerus size and foraging strategy in hummingbirds. Multivariate relationships between male humerus shape (PC1) and ecological and morphological predictors across 78 hummingbird species. Boxplots show group-level differences for categorical predictors. Scatterplots for continuous predictors show the estimated regression fit. Humerus shape varies with humerus size and foraging strategy, whereas other predictors show weak or non-significant associations. Silhouettes illustrate shape change along the PC1 axis. Body mass was analyzed as the natural logarithm of mass in grams, and geographic range size as the natural logarithm of area in km². Humerus size is derived from landmark coordinates, in relative (coordinate) units.

**Table 1.**
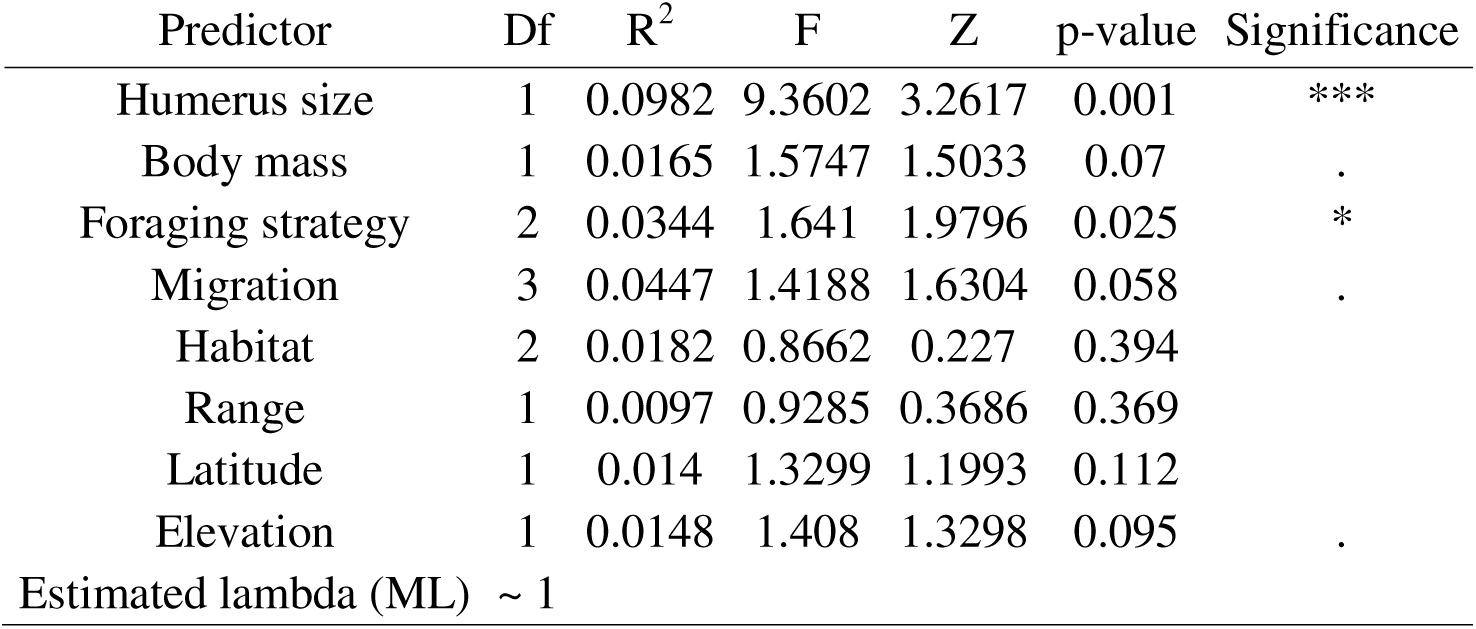
Humerus size is the primary predictor of humerus shape variation, with secondary effects of foraging strategy and migration. Results of multivariate PGLS models testing ecological and morphological predictors of male humerus shape across hummingbird species. For each predictor, degrees of freedom (Df), partial R^2^, F- and Z-statistics, and p-values are shown. Phylogenetic signal was strong, with maximum-likelihood, *λ* ≈ 1. Significance codes: *** p ≤ 0.001, * p ≤ 0.05, . p ≤ 0.1.

Across all predictors, effect sizes were modest, and the final model explained a limited fraction of total multivariate variation in humeral shape (ca. 25%), with size-related effects accounting for the largest share. Sensitivity analyses across values of Pagel’s λ showed that the effects of humerus size and foraging strategy were consistent (Table S7). The effects of humerus size and foraging strategy on humerus shape were also robust to the inclusion of an interaction between these two variables (Table S8).

### Morphospace differentiation by foraging strategy

PCA revealed substantial overlap in humeral morphospace among foraging strategies (Fig. 3A). The morphospaces of territorial and traplining species were largely contained within the broader morphospace occupied by opportunistic species, as illustrated by convex hulls. Projection of phylogenetic relationships onto the morphospace indicated some local clustering of related taxa, consistent with the detected phylogenetic signal, but no strong global structuring by clade.

**Figure 3.**
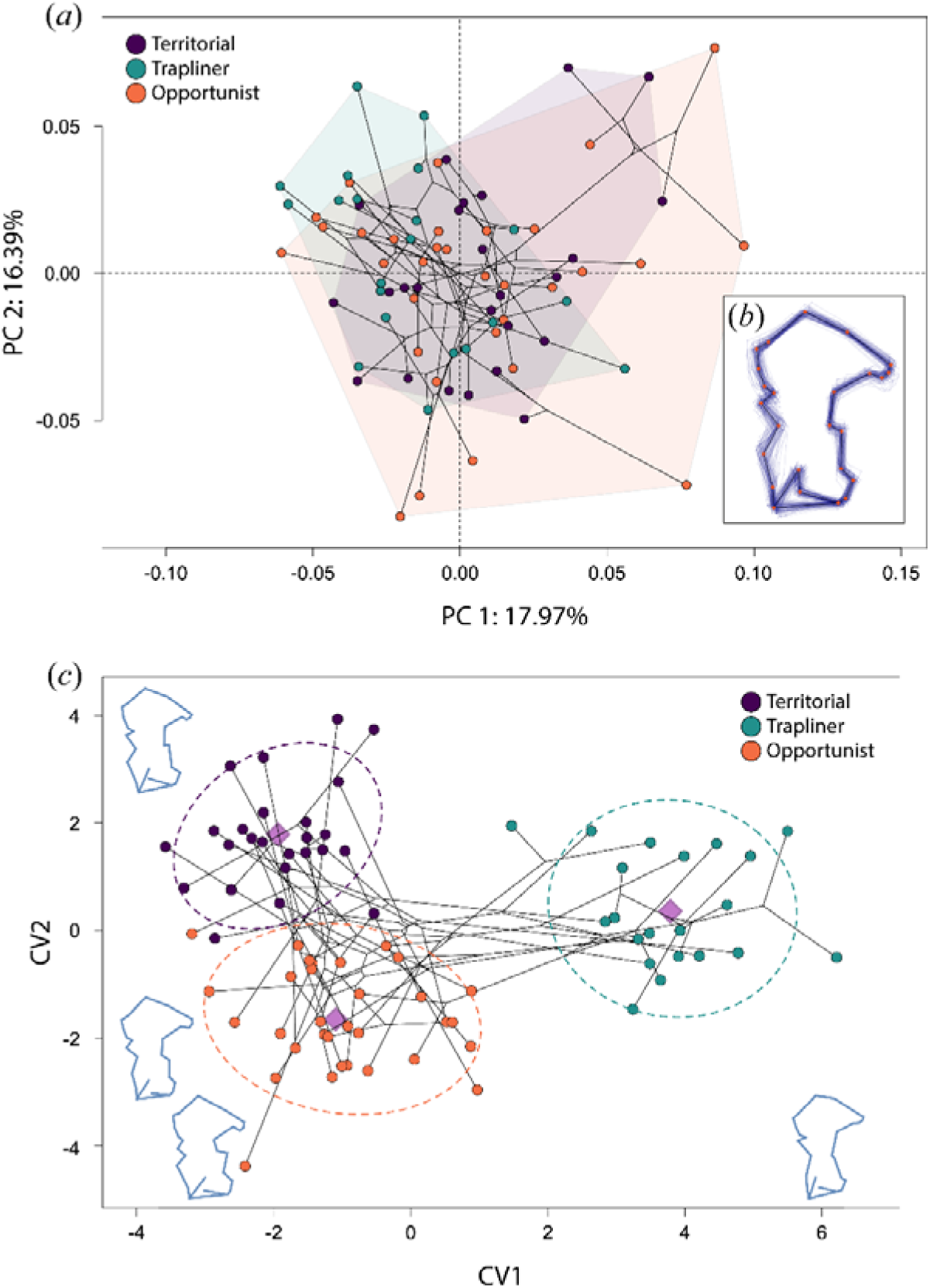
Hummingbird foraging strategies differ in humeral morphospace. Foraging strategies differ in their occupation of humeral morphospace, with territorial, traplining, and opportunistic species forming clusters in both unsupervised (PCA) and supervised (CVA) shape spaces. Each point represents a species (n = 78), colored by foraging strategy. (a) PCA of Procrustes-aligned humeral shapes, with PC1 and PC2 explaining 17.97% and 16.39% of variation, respectively. Shaded convex hulls denote group morphospaces, and thin black lines represent phylogenetic relationships projected into shape space. (b) Mean humeral shape variation across specimens visualized as a thin-plate spline deformation of the consensus configuration; red points denote landmarks. (c) Canonical Variate Analysis maximizing group separation based on foraging strategy. Dashed ellipses show 90% confidence regions, and purple diamonds mark group centroids. Shape outlines adjacent to the axes illustrate humeral changes associated with CV1 and CV2.

**Figure 4.**
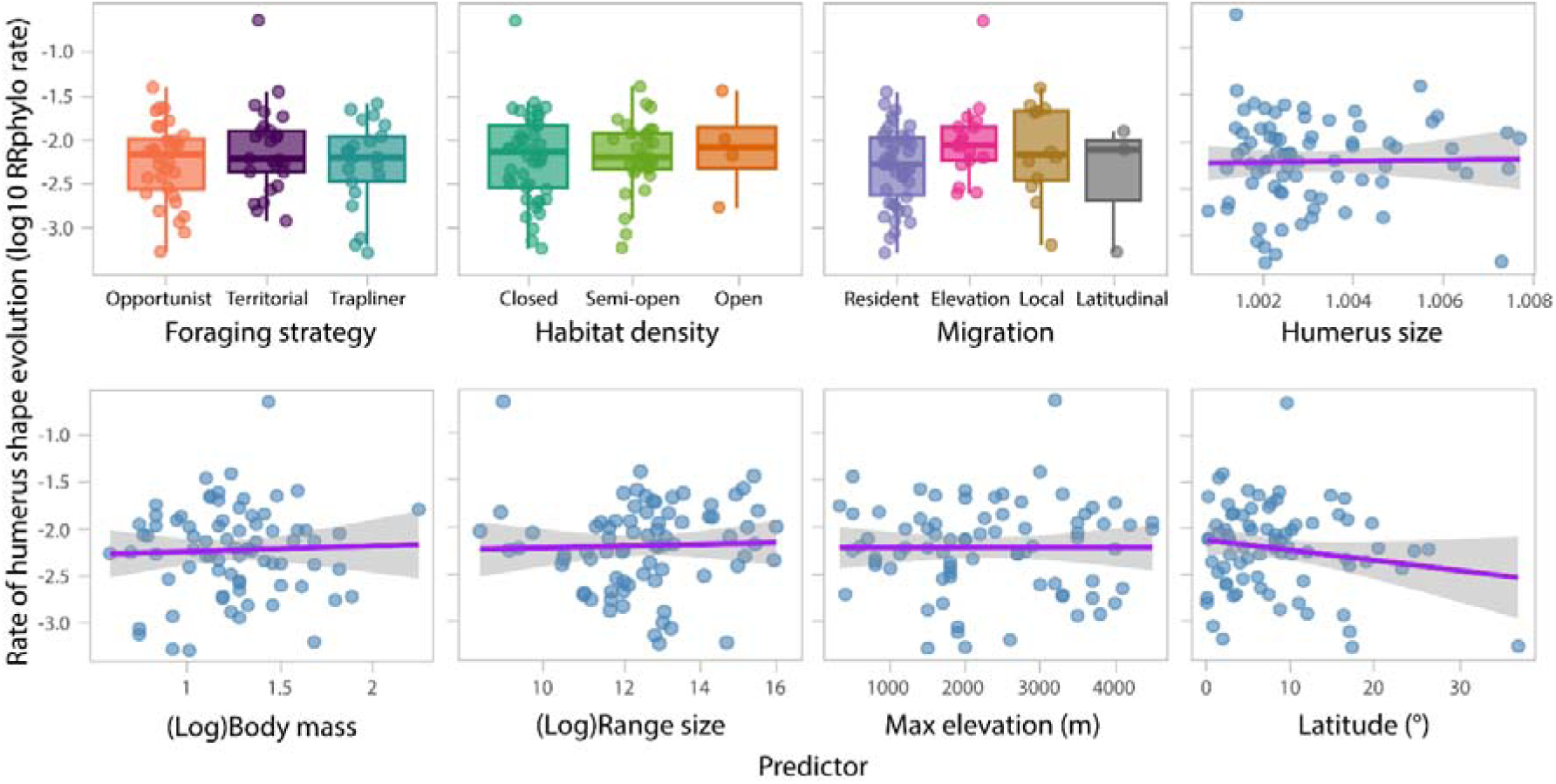
Evolutionary rates of hummingbird humerus shape show little association with ecological or morphological predictors. Relationships between estimated humerus shape evolutionary rates (log-transformed) and ecological and morphological predictors across 78 hummingbird species. Boxplots summarize group-level comparisons for categorical predictors, and scatterplots with fitted regression lines illustrate associations with continuous predictors. We do not detect associations between humerus shape evolutionary rates and any predictors. Body mass was analyzed as the natural logarithm of mass in grams, and geographic range size as the natural logarithm of area in km². Humerus size is derived from landmark coordinates, in relative (coordinate) units.

Canonical Variates Analysis further characterized differences among foraging strategies (Fig. 3C). The first canonical axis primarily separated traplining species from territorial and opportunistic taxa, whereas the second axis distinguished territorial from opportunistic species. However, cross-validated classification accuracy was low (42.3%), only slightly exceeding random expectations, and all strategies showed extensive overlap in canonical space. Traplining species exhibited higher correct classification (47.6%) than territorial and opportunistic species (36–43.7%). Consistent with these patterns, a permutational multivariate test on size-corrected shape data detected small but significant differences in mean humeral shape among foraging strategies (R² = 0.060, p = 0.001).

### Evolutionary rates of humeral shape

Estimates of evolutionary rates derived from *RRphylo* revealed a strongly right-skewed distribution, with most species exhibiting low rates of humeral shape evolution (median = 0.0063; interquartile range = 0.0030–0.0120, Fig. 1D). Only a small number of branches showed elevated rates, indicating isolated episodes of accelerated morphological change (Table S9). Phylogenetic signal in evolutionary rates was weak (λ ≈ 0.03) and not significantly different from zero (LR = 0.040, p = 0.840), indicating that closely related species did not evolve at more similar rates than expected under phylogenetic independence.

Group-wise comparisons detected no significant differences in evolutionary rates among foraging strategies (p = 0.138), habitat density categories (p = 0.503), or migratory classes (p = 0.953; Table 2). Similarly, univariate PGLS models showed no significant associations between evolutionary rates and continuous predictors, including body mass, humerus centroid size, geographic range size, elevation, or latitude (all p > 0.13; Table 2). These results indicate that variation in humeral evolutionary tempo is not systematically associated with these ecological or morphological predictors.

**Table 2.** Evolutionary rates of humerus shape do not differ significantly among ecological or morphological predictors. Group-wise rate variance analyses testing for differences in evolutionary rates of male humerus shape among categorical ecological predictors, and univariate PGLS models evaluating associations between evolutionary rates and continuous predictors. No predictor exhibited a statistically significant effect on humerus shape evolutionary rates.

| Group-wise rate variance analyses for categorical predictors |  |  |  |  |
| --- | --- | --- | --- | --- |
| Predictor | Observed Rate Ratio | Effect size | p-value |  |
| Foraging strategy | 2.5141 | 1.135 | 0.138 |  |
| Habitat | 2.0279 | -0.0196 | 0.503 |  |
| Migration | 1.6409 | -1.6145 | 0.953 |  |
| Univariate PGLS values |  |  |  |  |
| Predictor | Estimate | Std. Error | Lambda (ML) | p-value |
| Body mass | 0.0078 | 0.0092 | 0.0579 | 0.3996 |
| Humerus size | -1.1709 | 1.5639 | 0.0088 | 0.4563 |
| Range | -0.0026 | 0.0017 | 0.0327 | 0.1349 |
| Elevation | 0.000002 | 0.000003 | 0.0276 | 0.5455 |
| Latitude | -0.0001 | 0.0003 | 0.0217 | 0.8738 |

### Phenotypic modularity and integration

Phylogenetic partial least-squares analysis revealed strong integration between the proximal and distal regions of the humerus (r-PLS = 0.869, Z = 4.014, p = 0.001; Fig. 5). This indicates that morphological changes in one region are closely mirrored by changes in the other, consistent with coordinated evolution across the bone. The magnitude and significance of integration were robust to phylogenetic correction and did not depend on ecological or size-related predictors.

**Figure 5.**
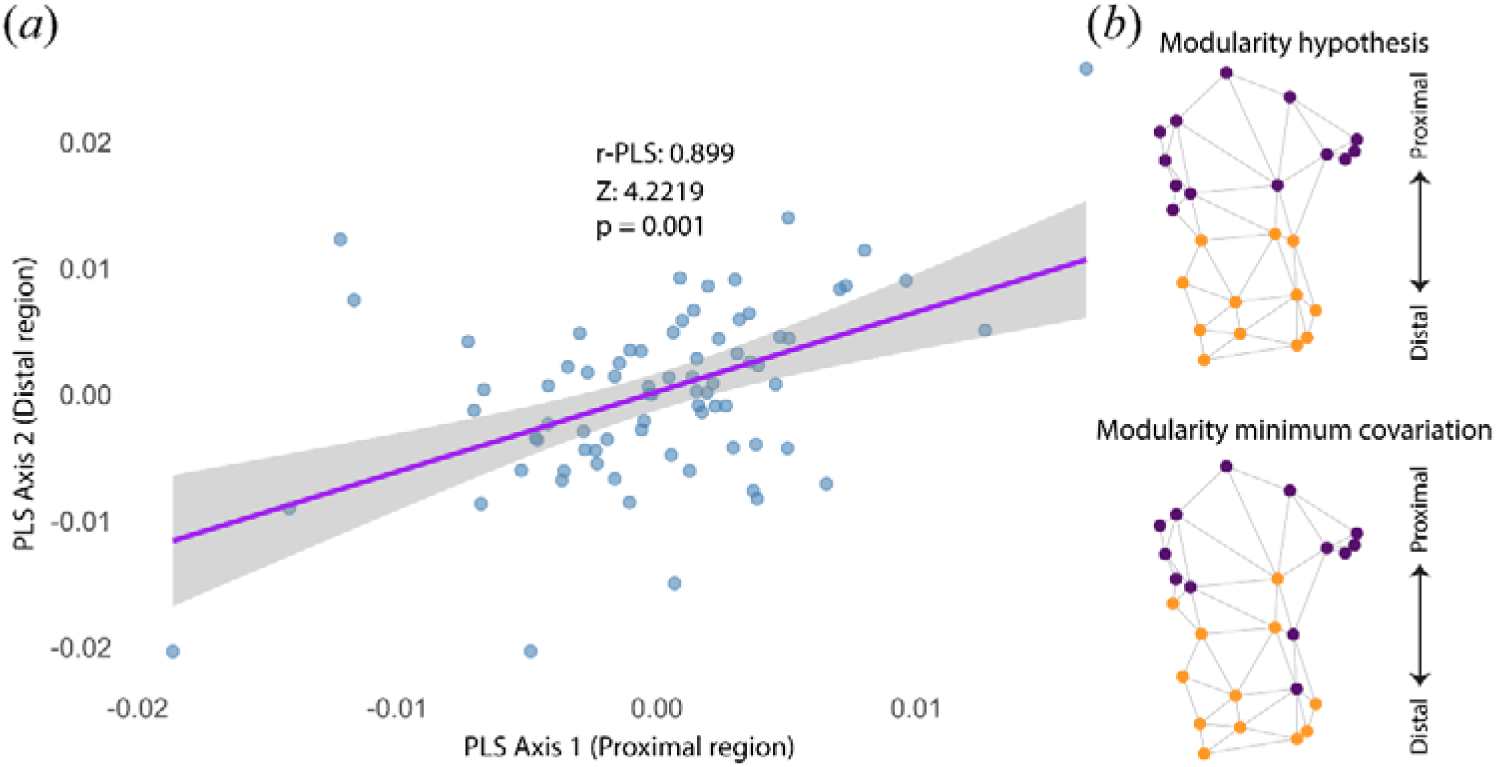
Proximal and distal regions of the hummingbird humerus are integrated, supporting a functional linkage rather than modular independence. The proximal and distal regions of the humerus exhibit morphological integration, indicating coordinated evolutionary changes rather than independent modular evolution. (a) Relationship between PLS1 scores of the proximal region and PLS2 scores of the distal region (r-PLS = 0.899, Z = 4.2219, p = 0.001). Each point represents a species (n = 78), and the shaded region shows the 95% confidence interval for this relationship. (b) The *a priori* modularity hypothesis (upper panel), representing the expected division of the humerus into proximal and distal functional units, contrasted with the data-driven minimum-covariation modular structure (lower panel). Nodes represent landmarks and edges represent the presence of covariation links among landmarks.

## Discussion

### Allometric scaling as the dominant driver of humeral shape evolution

Among all predictors evaluated, humerus centroid size showed the strongest and most consistent association with humeral shape, whereas body mass exerted only a marginal effect. This allometric signal indicates that increasing humeral size is associated with broader proximal epiphyses, increased mediolateral shaft robustness, and reduced curvature. Similar size-related reinforcement of proximal limb bones has been documented across volant birds and other flying vertebrates, where increases in cross-sectional dimensions enhance resistance to bending and torsional stresses [12,30,63]. These trends are consistent with general scaling principles linking skeletal strength to geometry rather than body mass per se [7,63].

Biologically, this pattern indicates that selection acting on flight performance in larger hummingbirds favors humeral morphologies that enhance joint stability and load-bearing capacity. Expanded articular surfaces increase muscle attachment area and force transmission at the shoulder, particularly for the pectoralis and supracoracoideus, whereas straighter and more robust diaphyses reduce bending moments generated during flapping and hovering [1,10,30]. Comparable relationships between bone robustness, joint morphology, and force transmission have been reported in avian forelimb mechanics [64,65]. This interpretation is especially informative given the unusual scaling of the hummingbird wing: distal components, particularly the manus and primary feathers, scale disproportionately with body mass, producing relatively long wings and low wing loading [8–10]. As a result, aerodynamic and inertial forces generated distally increase nonlinearly with size and must be transmitted proximally through the humerus. This contrast between elongation of distal wing elements and structural reinforcement of the humerus is consistent with previous work showing that elongation of distal elements contributes to increased aerodynamic efficiency [12,18,66]. Overall, these results show that allometric scaling imposes a dominant and predictable constraint on humeral evolution in hummingbirds, shaping mechanically robust configurations that stabilize force transmission during hovering and flapping flight across ecological contexts.

### Ecological effects as secondary modifiers of humeral shape

Although allometric scaling explains most of the variation in male humerus shape, our results suggest that foraging and competitive behavior act as a secondary modifier. Foraging strategy showed a statistically significant but modest association with humeral shape, with differences among species that use territorial, traplining, and opportunistic foraging behavior. These differences are modest, such that foraging categories overlap within a shared morphospace adapted to hovering flight, rather than discrete or mutually exclusive morphotypes. Principal component and canonical variate analyses also revealed that opportunistic hummingbird species span most of the variation occupied by territorial and traplining taxa. Although multivariate tests detected statistically significant differences in mean shape after accounting for size, these differences were small relative to overall variation. Similarly modest differentiation has been reported for other avian skeletal traits [67,68].

Despite their modest magnitude, associations between humerus shape and foraging behaviour were consistent with our predictions. Traplining species tend to exhibit relatively longer, more slender humeri, consistent with sustained forward flight over extended circuits, where elongation of the proximal wing segment may contribute to increased effective wingspan and reduced wingbeat frequency [12,13,66,69]. Territorial species tend to show shorter, thicker, more curved humeri; these features are expected to reduce shoulder moment of inertia, facilitating rapid angular acceleration and tight maneuvering during aggressive chases and vertical ascents [9,11,37,38].

Humeral elongation is not the sole route to increased wingspan or aerodynamic efficiency: similar outcomes can arise from elongation of distal wing elements, often with lower inertial costs [12,18]. Accordingly, we do not assume a one-to-one correspondence between humeral length and wing length; rather, humeral geometry likely interacts with distal skeletal and feather elements to shape overall wing planform and flight performance, making humeral elongation a plausible but not exclusive pathway contributing to strategy-associated differences. Our analyses also suggest that foraging effects may be size-dependent, rather than uniform. We found evidence for an interaction between foraging strategy and humerus centroid size (Table S8). These results suggest that foraging strategy may bias humeral shape evolution conditional on size, instead of acting as an independent driver of morphology.

Other ecological and geographic variables, including migratory behavior, habitat density, latitude, and elevation, showed marginal or no detectable associations with humeral shape, consistent with reports for other avian morphological traits once size and phylogeny are accounted for [25,70]. Habitat density and migratory categories are proxies for flight-space structure and movement ecology. Their lack of detectable associations with humerus shape suggests that they may primarily act on distal wing/feather components, or their effects may be overshadowed by size-related load-transmission constraints in the humerus.

The humerus—by virtue of its mechanically integrated, load-bearing role—may be buffered against many ecological pressures, with selection acting primarily within limits imposed by allometric scaling and structural integration [30,37]. Future studies using three-dimensional humeral models may reveal additional axes of shape variation—particularly related to articular surfaces and torsional geometry—not captured by the present two-dimensional approach, potentially uncovering stronger or complementary associations with ecological and functional predictors.

### Phylogenetic structure as background context

Across all analyses, humeral shape exhibited significant phylogenetic signal, indicating resemblance among related species. This pattern is typical of complex morphological traits and reflects shared ancestry, developmental pathways, and conserved biomechanical solutions [33,57]. Importantly, this phylogenetic structure provides the historical backdrop against which both shape variation and evolutionary rates are interpreted. By accounting for shared ancestry, size-dependent scaling and ecological effects can be evaluated as deviations from inherited morphological similarity, rather than artifacts of phylogenetic relatedness [39].

### Evolutionary rates: heterogeneity without ecological predictors

In contrast to patterns of shape variation, evolutionary rates of male humeral morphology showed no consistent association with ecological predictors. Rates were highly heterogeneous across the phylogeny, with most branches exhibiting low values and only a few showing elevated rates. Rate estimates also lacked detectable phylogenetic signal, and we did not detect any associations between evolutionary rates and any other species traits. Together, these patterns suggest that variation in evolutionary tempo arises idiosyncratically rather than being structured by shared ancestry [28,39]. Similar decoupling between morphological disparity and evolutionary tempo has been reported in other macroevolutionary studies, where strong constraints on form coexist with heterogeneous, lineage-specific rates of change [27,29].

These results caution against adaptive narratives that link particular ecological strategies to accelerated skeletal evolution without explicit support from evolutionary rate analyses. Although maximum elevation showed weak explanatory power in the multivariate models, individual lineages exhibited elevated evolutionary rates, including *Metallura phoebe*, a high-elevation species identified as a rate outlier (Table S9). Such cases suggest that localized increases in evolutionary rate may arise from recent, lineage-specific processes—such as shifts in functional integration, developmental constraints, or population-level dynamics—rather than reflecting a general effect of elevation on humeral evolution. Importantly, the presence of these outliers does not contradict the overall absence of a strong elevational signal, but instead highlights the distinction between global macroevolutionary trends and idiosyncratic evolutionary trajectories at the species level [29].

### Phenotypic integration as a key constraint on humeral evolution

We find significant phenotypic integration between proximal and distal regions of the hummingbird humerus. This finding indicates coordinated evolution, consistent with the shared role of the proximal and distal regions in force transmission during flight [1,30]. Such integration is expected in highly loaded locomotor structures, where changes in one region require compensatory adjustments elsewhere to preserve functional integrity [36,71].

This integration also provides an explanation for several observed patterns in our phylogenetic analysis. It helps explain why ecological effects on humeral shape are modest, as selection cannot act independently on isolated regions of the bone, and it indicates that evolutionary change in the humerus is channeled along constrained trajectories defined by internal structural coherence. Similar roles of integration in shaping evolutionary constraint have been documented across vertebrate skeletal systems [72,73]. Together, these results show that phenotypic integration simultaneously enables coordinated functional refinement and limits morphological disparity. Even when ecological pressures promote divergence in humeral shape, internal structural coherence remains evolutionarily conserved.

## Conclusion

Taken together, our results show that humeral evolution in hummingbirds is governed primarily by biomechanical scaling and internal structural integration, maintaining a skeletal configuration optimized for sustained hovering flight, with ecological factors introducing secondary, constrained modifications to shape. While behaviors such as foraging strategy leave detectable signatures on humeral morphology, these effects are modest relative to the dominant influence of size-dependent mechanics. Moreover, evolutionary rates of humeral change appear largely decoupled from ecological predictors, highlighting the importance of constraint and historical contingency in shaping macroevolutionary patterns. This integrative perspective underscores that, even within a lineage renowned for flight specialization, the evolution of skeletal form reflects a balance between universal biomechanical principles and the selective nuances of ecological diversification.

## Author contributions

**JCR:** Conceptualization, Methodology, Software, Formal analysis, Investigation, Data curation, Writing – Original draft, Writing – Review & Editing, Visualization. **RD:** Conceptualization, Validation, Investigation, Resources, Writing – Original draft, Writing – Review & Editing, Supervision, Funding acquisition. **CDC:** Conceptualization, Writing – Review & Editing, Supervision, Funding acquisition. **IB:** Investigation, Data curation, Writing – Review & Editing. **LM:** Investigation, Data curation. **ARG:** Conceptualization, Investigation, Writing – Review & Editing, Supervision.

## Acknowledgments

We thank Steven Cardiff and Nick Mason for their support during fieldwork and specimen access, as well as the Louisiana State University Museum of Natural Science for facilitating access to their ornithological collections.

## Data availability statement

Data and code for all analyses are available on Figshare at: https://figshare.com/s/2c012d7ba82c9eeda5ed

