## Supplementary material for "Scaling and foraging behavior drive the evolution of humeral shape in hummingbirds"

**Table S1. Specimens imaged for humeral landmarks, listed by clade, species, museum catalog number, and number of individuals.** For two of the species listed (*Calypte anna* and *Goethalsia bella*), the single male specimen did not pass quality-control criteria and hence those species were excluded from further analyses.

| Clade | Species | Male individuals | LSUMNS Catalog number | Female individuals | LSUMNS Catalog number |
| --- | --- | --- | --- | --- | --- |
| Bees | *Acestrura bombus* | 1 | 157294 | 0 | -- |
|  | *Archilochus alexandri* | 1 | 104000 | 1 | 164901 |
|  | *Calliphlox evelynae* | 1 | 86685 | 0 | -- |
|  | *Calypte anna* | 1 | 126295 | 1 | 126337 |
|  | *Selasphorus rufus* | 1 | 122561 | 0 | -- |
|  | *Selasphorus flammula* | 1 | 48518 | 1 | 48573 |
|  | *Thaumastura cora* | 1 | 157351 | 1 | 65301 |
| Brilliants | *Boissonneaua matthewsii* | 1 | 107081 | 0 | -- |
|  | *Coeligena coeligena* | 1 | 70210 | 2 | 129797 |
|  |  |  |  |  | 129799 |
|  | *Coeligena lutetiae* | 1 | 89783 | 0 | -- |
|  | *Eriocnemis aline* | 1 | 107088 | 1 | 107089 |
|  | *Eriocnemis luciani* | 2 | 99330 | 0 | -- |
|  |  |  | 99328 | 0 | -- |
|  | *Eriocnemis vestita* | 1 | 89829 | 1 | 89832 |
|  | *Heliodoxa branickii* | 1 | 107022 | 0 | -- |
|  | *Heliodoxa leadbeateri* | 1 | 172480 | 1 | 172481 |
|  | *Heliodoxa aurescens* | 2 | 107015 | 2 | 107014 |
|  |  |  | 118207 |  | 93889 |
|  | *Lafresnaya lafresnayi* | 1 | 75607 | 1 | 74850 |
|  | *Ocreatus underwoodii* | 1 | 89834 | 1 | 93902 |
|  | *Pterophanes cyanopterus* | 2 | 129794 | 1 | 79763 |
|  |  |  | 101268 |  |  |
|  | *Urosticte benjamini* | 1 | 157298 | 0 | -- |
| Coquettes | *Adelomyia melanogenys* | 1 | 83967 | 1 | 89749 |
|  | *Chalcostigma herrani* | 1 | 170439 | 1 | 170440 |
|  | *Chalcostigma ruficeps* | 1 | 70215 | 0 | -- |
|  | *Aglaiocercus kingii* | 1 | 118211 | 2 | 71576 |
|  |  |  |  |  | 97489 |
|  | *Chalcostigma stanleyi* | 2 | 79780 | 0 | -- |
|  |  |  | 93937 |  |  |
|  | *Heliangelus amethysticollis* | 1 | 101274 | 1 | 68833 |
|  | *Heliangelus exortis* | 1 | 160769 | 1 | 160770 |
|  | *Lesbia victoriae* | 1 | 93903 | 0 | -- |
|  | *Lophornis delattrei* | 1 | 171196 | 0 | -- |
|  | *Metallura phoebe* | 1 | 65300 | 0 | -- |
|  | *Metallura tyrianthina* | 1 | 74866 | 0 | -- |
|  | *Sappho sparganurus* | 1 | 101275 | 1 | 95373 |
| Emeralds | *Amazilia amazilia* | 1 | 157373 | 1 | 93882 |
|  | *Amazilia candida* | 2 | 31873 | 2 | 31875 |
|  |  |  | 31871 |  | 31872 |
|  | *Amazilia chionogaster* | 3 | 125751 | 1 | 172508 |
|  |  |  | 125749 |  |  |
|  |  |  | 125748 |  |  |
|  | *Amazilia lactea* | 1 | 157368 | 3 | 157367 |
|  |  |  |  |  | 157366 |
|  |  |  |  |  | 157365 |
|  | *Amazilia yucatanensis* | 1 | 168115 | 0 | -- |
|  | *Amazilia cyanocephala* | 1 | 31880 | 1 | 31879 |
|  | *Amazilia saucerottei* | 1 | 32032 | 0 | -- |
|  | *Campylopterus villaviscensio* | 1 | 160856 | 0 | -- |
|  | *Chalybura buffonii* | 1 | 108747 | 1 | 93884 |
|  | *Chlorostilbon lucidus* | 1 | 125732 | 0 | -- |
|  | *Chlorestes notata* | 1 | 118340 | 0 | -- |
|  | *Chlorostilbon canivetii* | 1 | 31868 | 0 | -- |
|  | *Chrysuronia oenone* | 4 | 70207 | 0 | -- |
|  |  |  | 118206 |  |  |
|  |  |  | 70206 |  |  |
|  |  |  | 125743 |  |  |
|  | *Damophila julie* | 1 | 160759 | 1 | 160762 |
|  | *Elvira chionura* | 1 | 50762 | 0 | -- |
|  | *Eupherusa eximia* | 1 | 163857 | 0 | -- |
|  | *Goethalsia bella* | 1 | 108746 | 0 | -- |
|  | *Hylocharis chrysura* | 1 | 125742 | 0 | -- |
|  | *Hylocharis leucotis* | 1 | 14766 | 1 | 31870 |
|  | *Klais guimeti* | 1 | 50765 | 1 | 86440 |
|  | *Leucippus baeri* | 1 | 93880 | 0 | -- |
|  | *Leucippus chlorocercus* | 1 | 121031 | 1 | 121030 |
|  | *Leucippus taczanowskii* | 1 | 93881 | 1 | 81208 |
|  | *Orthorhyncus cristatus* | 2 | 150467 | 0 | -- |
|  |  |  | 150466 |  |  |
|  | *Thalurania furcata* | 1 | 50758 | 1 | 118343 |
|  | *Thalurania colombica* | 1 | 108743 | 2 | 108745 |
|  |  |  |  |  | 48568 |
| Hermits | *Eutoxeres condamini* | 1 | 74844 | 0 | -- |
|  | *Glaucis aeneus* | 1 | 50745 | 3 | 77733 |
|  |  |  |  |  | 48754 |
|  |  |  |  |  | 48563 |
|  | *Glaucis hirsutus* | 2 | 118576 | 0 | -- |
|  |  |  | 118577 |  |  |
|  | *Phaethornis bourcieri* | 3 | 121013 | 1 | 121012 |
|  |  |  | 111170 |  |  |
|  |  |  | 111169 |  |  |
|  | *Phaethornis hispidus* | 1 | 111160 | 1 | 111159 |
|  | *Phaethornis koepckeae* | 2 | 118193 | 1 | 118190 |
|  |  |  | 118192 |  |  |
|  | *Phaethornis pretrei* | 1 | 125715 | 0 | -- |
|  | *Phaethornis striigularis* | 1 | 108740 | 1 | 31864 |
|  | *Phaethornis subochraceus* | 1 | 125717 | 0 | -- |
|  | *Phaethornis superciliosus* | 1 | 111157 | 2 | 111153 |
|  |  |  |  |  | 156879 |
|  | *Phaethornis yaruqui* | 1 | 68731 | 0 | -- |
|  | *Phaethornis philippii* | 2 | 118338 | 1 | 118337 |
|  |  |  | 118332 |  |  |
|  | *Threnetes leucurus* | 1 | 89720 | 1 | 89905 |
|  | *Threnetes ruckeri* | 1 | 77735 | 1 | 62775 |
| Mangoes | *Anthracothorax dominicus* | 1 | 150456 | 0 | -- |
|  | *Colibri cyanotus* | 1 | 83965 | 3 | 70204 |
|  |  |  |  |  | 83964 |
|  |  |  |  |  | 81209 |
|  | *Doryfera johannae* | 1 | 118178 | 2 | 118179 |
|  |  |  |  |  | 89714 |
|  | *Doryfera ludovicae* | 1 | 89719 | 0 | -- |
|  | *Polytmus guainumbi* | 1 | 125744 | 0 | -- |
|  | *Schistes geoffroyi* | 2 | 89850 | 1 | 68832 |
|  |  |  | 89849 |  |  |
| Mtn.Gems | *Heliomaster longirostris* | 1 | 153956 | 0 | -- |
|  | *Panterpe insignis* | 2 | 48570 | 0 | -- |
|  |  |  | 64845 |  |  |

**Table S2. Anatomical definition of humeral landmarks used in geometric morphometric analyses.** List of 25 landmarks categorized by anatomical region (proximal epiphysis, diaphysis, distal epiphysis) with descriptions of their osteological positions.

| Bone region | Landmark | Description |
| --- | --- | --- |
| Proximal epiphysis | 1 | Proximal end of ventral tubercle |
|  | 2 | Ventral end of the head notch |
|  | 3 | Most proximal end of the humeral head |
|  | 4 | More depressed area at the junction between the deltopectoral ridge and the humeral head |
|  | 5 | Proximal end of the deltopectoral crest intersecting with the humeral head |
|  | 6 | Medial depression of the deltopectoral crest |
|  | 7 | Most distal end of the deltopectoral crest |
|  | 8 | Origin of the deltopectoral crest in the diaphysis |
|  | 22 | Point of greatest convexity in joint between the bicipital crest and the diaphysis |
|  | 23 | Most dorsal-proximal end of the ventral tubercule |
|  | 24 | Most distal end of the ventral tubercule |
|  | 25 | Most ventral end of the ventral tubercule |
| Diaphysis | 9 | Dorsal end of the extensor metacarpi radialis process |
|  | 10 | Distal origin of the extensor metacarpi radialis process |
|  | 20 | Ventral-proximal end of the medio-ventral tubercule |
|  | 21 | Proximal origin of the medio-ventral tubercule |
| Distal epiphysis | 11 | Dorsal end of the tubercule of tensor propatagialis, pars brevis |
|  | 12 | Proximal origin of the tubercule of tensor propatagialis, pars brevis |
|  | 13 | Most dorsal-distal end of the dorsal condyle |
|  | 14 | Most ventral-proximal end of the dorsal condyle |
|  | 15 | Most dorsal-proximal end of the ventral condyle |
|  | 16 | Most ventral-distal end of the ventral condyle |
|  | 17 | Distal origin of the ventral epicondyle |
|  | 18 | Most ventral end of the ventral epicondyle |
|  | 19 | Proximal origin of the ventral epicondyle |

**Documentation of foraging strategy assignments used in this study**

**Operational definitions of foraging strategies**

We defined territorial foraging species as those described as predominantly defending spatially fixed floral resources, including repeated aggressive interactions (chasing, displacement, or dominance displays) that exclude heterospecific or conspecific competitors from feeding patches. Species classified as territorial are consistently described as maintaining feeding territories, even when floral density is moderate to high. We defined traplining species as those described as using repeated visitation of spatially dispersed flowers along relatively stable foraging routes, with little or no territorial defense of floral resources. Traplining species are typically described as avoiding aggressive interactions and relying on circuit-based foraging across multiple plants or patches. We defined opportunistic species as those described as using flexible or context-dependent foraging behavior, including both territory defense and non-territorial foraging, depending on resource abundance, competitive environment, or season. Species described as facultatively territorial, or exhibiting mixed strategies across populations were classified as opportunistic.

**Assignment rules and handling of ambiguous cases**

We assigned species to the foraging category that best represents their dominant strategy based on the sources outlined in Table S3 below. When descriptions differed among sources or populations, priority was given to (i) primary behavioral studies conducted during breeding, followed by (ii) regional species accounts and comparative syntheses. Species exhibiting persistent mixed strategies without clear predominance were classified as opportunistic.

**Limitations**

We recognize that classifying species into discrete foraging categories simplifies a continuous behavioral spectrum in hummingbirds, and that species may shift strategies across seasons or ecological conditions. As a result, categorical assignments cannot capture all behavioral nuance. Nonetheless, these broad foraging strategies reflect recurrent differences in resource use that are expected to impose distinct selective pressures on morphology over evolutionary timescales, making them a useful comparative framework when interpreted cautiously.

**Table S3. Foraging strategies for hummingbird species in this study.** For each species, the table provides the original assignment reported by Rombaut et al., (2022), the category used in our analyses, and additional supporting notes used in our designation.

| Species | Foraging strategy (Rombault et al, 2022) | Foraging strategy assigned for this study | Additional notes used for ambiguous cases |
| --- | --- | --- | --- |
| *Acestrura bombus* | -(Not included)- | Opportunist | "Foraging strategy presumably similar to C. mulsant" |
| *Archilochus alexandri* | Opportunist | Opportunist | "Uses a multitude of different plant species throughout its large geographic range and within its many varied habitats" |
| *Calliphlox evelynae* | Territorial | Territorial |  |
| *Calypte anna* | Territorial | Territorial |  |
| *Selasphorus rufus* | Territorial | Territorial |  |
| *Selasphorus flammula* | Opportunist | Territorial | "In the non-breeding season, both sexes may defend territories around certain patches of small flowers... Male and sometimes female may defend feeding territories at large clumps of flowers, especially outside the breeding season, if not excluded by larger, more dominant species" |
| *Thaumastura cora* | Territorial | Territorial |  |
| *Boissonneaua matthewsii* | -(Not included)- | Territorial |  |
| *Coeligena coeligena* | Trapliner | Opportunist | "Feeds by trap-lining in lower to middle strata within and at edge of forest. Seldom seen in canopy of flowering trees where occasionally defends feeding territories." |
| *Coeligena lutetiae* | Opportunist | Trapliner | "Trap-liner, foraging at lower level at forest edges" |
| *Eriocnemis aline* | -(Not included)- | Opportunist | "Mainly takes nectar from flowers at 1–3 m above ground inside dense vegetation" |
| *Eriocnemis luciani* | Territorial | Territorial |  |
| *Eriocnemis vestita* | Territorial | Territorial |  |
| *Heliodoxa aurescens* | Trapliner | Trapliner |  |
| *Heliodoxa branickii* | -(Not included)- | Opportunist | "forages in the understory or along the edge of forests along the east slope of the Andes of Peru and northern Bolivia, especially on outlying ridges...very little information on the flowering plants visited by this hummingbird" |
| *Heliodoxa leadbeateri* | -(Not included)- | Trapliner | " It forages at flowers in the mid-story … Takes nectar mainly at flowers in lower to middle strata (1–10 m) inside forest or near small openings at forest edge. Usually forages alone and does not gather in groups at flowering trees" |
| *Lafresnaya lafresnayi* | Territorial | Territorial |  |
| *Ocreatus underwoodii* | -(Not included)- | Territorial | "Males establish feeding territories in which conspecific females and young males are allowed to feed, though their nectar-feeding activities were often interrupted by the territory owner…" |
| *Pterophanes cyanopterus* | Opportunist | Opportunist | "Territorial video , but may occasionally also trap-line flowers. Insects are caught by hawking." |
| *Urosticte benjamini* | -(Not included)- | Opportunist | "They typically forage by themselves, taking nectar from small flowers...Nectar of flowering Ericaceae, Fabaceae (Inga), Rubiaceae and bromeliads" |
| *Adelomyia melanogenys* | -(Not included)- | Trapliner | "feed at scattered flowers" |
| *Aglaiocercus kingii* | Opportunist | Opportunist |  |
| *Chalcostigma herrani* | Territorial | Territorial |  |
| *Chalcostigma ruficeps* | Territorial | Territorial |  |
| *Chalcostigma stanleyi* | Territorial | Territorial |  |
| *Heliangelus amethysticollis* | Territorial | Territorial |  |
| *Heliangelus exortis* | Territorial | Territorial |  |
| *Lesbia victoriae* | -(Not included)- | Opportunist | "...forages in montane scrub, in gardens, and other semiopen habitats...Forages for nectar at middle to high strata of flowering Bignoniaceae, Fabaceae, Gesneriaceae, Leguminosaceae, Puya and introduced Eucalyptus trees" NOTE: "Differences in display and behaviour do not suggest particularly close relationship with L. nuna" |
| *Lophornis delattrei* | -(Not included)- | Trapliner | No details on foraging strategy on BOW. Assigned using near species, unclear |
| *Metallura phoebe* | Territorial | Territorial |  |
| *Metallura tyrianthina* | Territorial | Territorial |  |
| *Sappho sparganurus* | Territorial | Territorial |  |
| *Amazilia amazilia* | Territorial | Territorial |  |
| *Amazilia candida* | Trapliner | Trapliner |  |
| *Amazilia chionogaster* | -(Not included)- | Opportunist | "Feeds on nectar of various plants of different families, such as Leguminosae, Vochysiaceae, Malvaceae, Lorantaceae, Bignoniaceae, Bombacaceae, Passifloraceae, Musaceae" |
| *Amazilia cyanocephala* | Opportunist | Opportunist | "There is little information on the diet… usually is solitary, as is typical of hummingbirds, but may aggregate at flowering trees" |
| *Amazilia lactea* | Territorial | Territorial |  |
| *Amazilia saucerottei* | Territorial | Territorial |  |
| *Amazilia yucatanensis* | Territorial | Territorial |  |
| *Campylopterus villaviscensio* | -(Not included)- | Opportunist | "Feeds on nectar, presumably of flowering Heliconia, bromeliads and other plants" |
| *Chalybura buffonii* | Opportunist | Opportunist |  |
| *Chlorostilbon lucidus* | Trapliner | Trapliner | "Feeds by trap-lining. Takes insect honeydew of Coccidae at Mimosa bracaatinga. Insects are caught in the air by hawking." |
| *Chlorostilbon canivetii* | Trapliner | Trapliner |  |
| *Chlorestes notata* | Territorial | Territorial |  |
| *Chrysuronia oenone* | Opportunist | Opportunist | "female more often trap-lines dispersed flowers in forest, male occasionally defends territories at flowers when more aggressive species like Thalurania furcata are absent" |
| *Damophila julie* | Territorial | Territorial |  |
| *Elvira chionura* | -(Not included)- | Opportunist | "Nectar of flowering shrubs and vines along forest borders and within forest." |
| *Eupherusa eximia* | Opportunist | Territorial |  |
| *Goethalsia bella* | -(Not included)- | Opportunist | "Pirre Hummingbird usually forages at flowers of shrubs or small trees at low or mid levels inside the forest... At one site it was reported to forage almost exclusively at red and blue flowers on a forest understory shrub" |
| *Hylocharis chrysura* | Opportunist | Opportunist |  |
| *Hylocharis leucotis* | Opportunist | Opportunist | Many plant species listed as food sources in botw |
| *Klais guimeti* | Opportunist | Territorial |  |
| *Leucippus baeri* | -(Not included)- | Trapliner | "Tumbes Hummingbird feeds on nectar from dry-tolerant plants like cacti…there is little information on the flowering plants visited by this hummingbird...There are no published data on territorial defense..." |
| *Leucippus chlorocercus* | -(Not included)- | Trapliner | "Forages for nectar at species of diverse plant families, including Leguminosae, Lorantaceae, Vochysiaceae, Bro­me­liaceae, Rutaceae, Bombacaceae, Malva­ceae, Rubiaceae, Myrtaceae, Passiflora­ceae" |
| *Leucippus taczanowskii* | -(Not included)- | Trapliner | "occupies arid scrub or the edges of dry forests, and feeds on nectar of plants like Agave or banana… little is known about the natural history of this species...visiting flowering Inga" |
| *Orthorhyncus cristatus* | -(Not included)- | Opportunist | "feeds on nectar from a variety of flowers...Nectar of flowering shrubs (Lantana, Euphorbia), vines and from lower parts of hedges and trees (Hibiscus, Bauhinia, Tabebuia, Delonix)" |
| *Thalurania colombica* | Territorial | Territorial |  |
| *Thalurania furcata* | Opportunist | Opportunist | "While males sometimes defend patches of flowers and act aggressively, females tend to be less territorial and either trapline or steal nectar from the floral territories of other hummingbirds" |
| *Heliomaster longirostris* | Opportunist | Opportunist |  |
| *Panterpe insignis* | Territorial | Territorial |  |
| *Eutoxeres condamini* | Trapliner | Trapliner |  |
| *Glaucis aeneus* | -(Not included)- | Opportunist | "Nectar, e.g. of Heliconia species", unclear |
| *Glaucis hirsutus* | -(Not included)- | Opportunist | "Nectarof Heliconia (e.g. H. bihai) (1), Centropogon (e.g. C. surinamensis) (1), Pachystachys, Passiflora, Trichanthera and Costus species, as well as small arthropods" |
| *Phaethornis bourcieri* | Trapliner | Trapliner |  |
| *Phaethornis hispidus* | Trapliner | Trapliner |  |
| *Phaethornis koepckeae* | Trapliner | Trapliner |  |
| *Phaethornis philippii* | Trapliner | Trapliner |  |
| *Phaethornis pretrei* | Trapliner | Trapliner |  |
| *Phaethornis striigularis* | Trapliner | Trapliner |  |
| *Phaethornis subochraceus* | -(Not included)- | Trapliner | based on: https://birdsofbolivia.org/species-fact-sheets-2/hummingbirds-picaflores/phaethornis-subochraceus/ |
| *Phaethornis superciliosus* | Trapliner | Trapliner |  |
| *Phaethornis yaruqui* | Trapliner | Trapliner |  |
| *Threnetes leucurus* | Trapliner | Trapliner |  |
| *Threnetes ruckeri* | Trapliner | Trapliner |  |
| *Anthracothorax dominicus* | Territorial | Territorial |  |
| *Colibri cyanotus* | Opportunist | Opportunist | "Lesser Violetear is flexible in its foraging behavior, depending upon flower density and the presence of competing species" |
| *Doryfera johannae* | Trapliner | Trapliner |  |
| *Doryfera ludovicae* | Trapliner | Trapliner |  |
| *Polytmus guainumbi* | -(Not included)- | Opportunist | "Nectar of flowering garden plants such as Lagerstroemia, low-growing shrubs like Lagerstroemia, Russelia equisetiformis and Calliandra surinamensis, clumps of Heliconia, Leguminosae, Malvaceae, Rubiaceae or Verbenaceae. In Argentina, observed feeding at the following plants and trees: Ceiba speciosa, Handroanthus heptaphyllus and Eriobotrya japonica" |
| *Schistes geoffroyi* | -(Not included)- | Opportunist | "feed from tubular flowers and often rob nectar by piercing the base of flowers without actually achieving pollination...Nectar of flowering shrubs, vines and small trees, e.g. Centropogon, Fuschia, Psammisia, Cavendishia, Palicourea, Besleria" |

**Table S4. Relative explanatory power of ecological, flight, and extended predictor sets for humerus shape variation.** Comparison of partial $R^{2}$ values from three multivariate PGLS models evaluating different predictor sets. We present a model including ecological traits, a model including flight-related traits, and an extended traits model combining ecological, flight, and morphological predictors. Partial $R^{2}$, p-values, and significance levels are reported for each predictor. In the extended traits model, humerus size shows the highest explanatory power, whereas other predictors exhibit weak or marginal effects.

| Model | Predictor | R^2^ | p-value | Significance |
| --- | --- | --- | --- | --- |
| Ecological traits | Foraging strategy | 0.0380 | 0.137 |  |
|  | Habitat | 0.0199 | 0.546 |  |
| Flight traits | Elevation | 0.0336 | 0.015 | * |
|  | Body mass | 0.0475 | 0.018 | * |
|  | Migration | 0.0465 | 0.169 |  |
| Extended traits | Foraging strategy | 0.0344 | 0.060 | . |
|  | Elevation | 0.0148 | 0.156 |  |
|  | Body mass | 0.0165 | 0.119 |  |
|  | Humerus size | 0.0982 | 0.005 | ** |
|  | Range | 0.0097 | 0.491 |  |
|  | Latitude | 0.0140 | 0.190 |  |
|  | Migration | 0.0447 | 0.105 |  |
|  | Habitat | 0.0182 | 0.584 |  |

**Table S5. Modeled predictors show low collinearity and meet assumptions for multivariate analyses.** Pearson and Spearman correlation matrices among continuous predictors included in our PGLS analyses, along with variance inflation factor (VIF) diagnostics. Correlation coefficients are uniformly low, and adjusted VIF values indicate no problematic collinearity.

| Pearson correlation matrix | | | | | |
| --- | --- | --- | --- | --- | --- |
|  | Humerus size | Body mass | Range | Latitude | Elevation |
| Humerus size | 1.000 | 0.064 | 0.015 | 0.170 | 0.008 |
| Body mass | 0.064 | 1.000 | 0.170 | -0.339 | 0.146 |
| Range | 0.015 | 0.170 | 1.000 | 0.059 | -0.295 |
| Latitude | 0.170 | -0.339 | 0.059 | 1.000 | -0.128 |
| Elevation | 0.008 | 0.146 | -0.295 | -0.128 | 1.000 |
| Spearman correlation matrix | | | | | |
|  | Humerus size | Body mass | Range | Latitude | Elevation |
| Humerus size | 1.000 | 0.178 | -0.014 | 0.012 | 0.052 |
| Body mass | 0.178 | 1.000 | 0.051 | -0.346 | 0.163 |
| Range | -0.014 | 0.051 | 1.000 | 0.017 | -0.317 |
| Latitude | 0.012 | -0.346 | 0.017 | 1.000 | -0.147 |
| Elevation | 0.052 | 0.163 | -0.317 | -0.147 | 1.000 |
| Variance inflation factor | | | | | |
|  | GVIF | Df | GVIF^(1/(2*Df)) |  |  |
| Humerus size | 1.187579 | 1 | 1.089761 |  |  |
| Body mass | 1.271138 | 1 | 1.127448 |  |  |
| Foraging | 1.883646 | 2 | 1.171520 |  |  |
| Nigration | 1.957897 | 3 | 1.118489 |  |  |
| Habitat | 1.631981 | 2 | 1.130261 |  |  |
| Range | 1.359011 | 1 | 1.165766 |  |  |
| Latitude | 1.881517 | 1 | 1.371684 |  |  |
| Elevation | 1.676100 | 1 | 1.294643 |  |  |

**Table S6. Type I p-values vary with predictor order, indicating order-dependent inference.** Evaluation of Type I (sequential) multivariate PGLS results under alternative orders of predictor entry. Each row represents a different ordering of predictors, with corresponding p-values shown for each sequential position. Statistical significance under Type I sums of squares depends on predictor order, motivating the use of order-invariant analyses in the main text.

| Model order | p 1 | p 2 | p 3 | p 4 | p 5 | p 6 | p 7 | p 8 |
| --- | --- | --- | --- | --- | --- | --- | --- | --- |
| Hum Size + Forag. + Elev. + Hab. + Range + Lat. + Mig. + B mass | 0.002 | 0.07 | 0.087 | 0.426 | 0.185 | 0.162 | 0.081 | 0.119 |
| Hum Size + Hab. + Lat. + B mass + Mig. + Elev. + Range + Forag. | 0.002 | 0.551 | 0.063 | 0.056 | 0.115 | 0.18 | 0.463 | 0.06 |
| Hum Size + Range + Lat. + Mig. + Elev. + B mass + Forag. + Hab. | 0.002 | 0.269 | 0.091 | 0.075 | 0.111 | 0.118 | 0.067 | 0.584 |
| Hum Size + Range + Lat. + Mig. + Forag. + B mass + Elev. + Hab. | 0.002 | 0.269 | 0.091 | 0.075 | 0.067 | 0.083 | 0.132 | 0.584 |
| B mass + Lat. + Elev. + Forag. + Range + Hum Size + Hab. + Mig. | 0.02 | 0.042 | 0.027 | 0.087 | 0.284 | 0.004 | 0.448 | 0.105 |
| B mass + Lat. + Mig. + Range + Elev. + Forag. + Hum Size + Hab. | 0.02 | 0.042 | 0.074 | 0.407 | 0.027 | 0.069 | 0.004 | 0.584 |
| B mass + Elev. + Hum Size + Mig. + Lat. + Range + Hab. + Forag. | 0.02 | 0.013 | 0.004 | 0.056 | 0.281 | 0.502 | 0.602 | 0.06 |
| B mass + Mig. + Range + Forag. + Hum Size + Hab. + Lat. + Elev. | 0.02 | 0.123 | 0.371 | 0.103 | 0.008 | 0.565 | 0.144 | 0.156 |
| Elev. + Hab. + Mig. + Forag. + Range + B mass + Hum Size + Lat. | 0.031 | 0.437 | 0.1 | 0.063 | 0.214 | 0.018 | 0.008 | 0.19 |
| Elev. + Hab. + Range + Mig. + Hum Size + Forag. + Lat. + B mass | 0.031 | 0.437 | 0.121 | 0.111 | 0.005 | 0.081 | 0.128 | 0.119 |
| Elev. + Hum Size + Mig. + Range + Forag. + Hab. + Lat. + B mass | 0.031 | 0.003 | 0.053 | 0.544 | 0.087 | 0.486 | 0.128 | 0.119 |
| Elev. + Forag. + Lat. + Range + Hab. + Mig. + B mass + Hum Size | 0.031 | 0.085 | 0.047 | 0.376 | 0.337 | 0.067 | 0.018 | 0.005 |
| Lat. + Elev. + B mass + Forag. + Hab. + Hum Size + Range + Mig. | 0.047 | 0.052 | 0.02 | 0.087 | 0.415 | 0.007 | 0.352 | 0.105 |
| Lat. + Forag. + B mass + Elev. + Mig. + Range + Hab. + Hum Size | 0.047 | 0.059 | 0.02 | 0.033 | 0.06 | 0.546 | 0.481 | 0.005 |
| Lat. + Hab. + Elev. + Range + B mass + Hum Size + Mig. + Forag. | 0.047 | 0.474 | 0.057 | 0.281 | 0.02 | 0.002 | 0.133 | 0.06 |
| Lat. + B mass + Hab. + Hum Size + Mig. + Range + Forag. + Elev. | 0.047 | 0.021 | 0.551 | 0.002 | 0.115 | 0.47 | 0.061 | 0.156 |
| Forag. + B mass + Hum Size + Lat. + Mig. + Range + Hab. + Elev. | 0.094 | 0.022 | 0.005 | 0.098 | 0.084 | 0.545 | 0.517 | 0.156 |
| Forag. + Hab. + Elev. + Range + B mass + Hum Size + Lat. + Mig. | 0.094 | 0.336 | 0.026 | 0.114 | 0.02 | 0.007 | 0.228 | 0.105 |
| Forag. + B mass + Hab. + Lat. + Hum Size + Elev. + Mig. + Range | 0.094 | 0.022 | 0.424 | 0.036 | 0.005 | 0.193 | 0.087 | 0.491 |
| Forag. + Hum Size + Range + B mass + Mig. + Hab. + Lat. + Elev. | 0.094 | 0.006 | 0.238 | 0.027 | 0.104 | 0.565 | 0.144 | 0.156 |
| Range + Lat. + Hum Size + Mig. + Hab. + Elev. + Forag. + B mass | 0.105 | 0.061 | 0.001 | 0.075 | 0.533 | 0.121 | 0.057 | 0.119 |
| Range + Lat. + Mig. + Elev. + Hab. + Forag. + B mass + Hum Size | 0.105 | 0.061 | 0.078 | 0.044 | 0.489 | 0.043 | 0.018 | 0.005 |
| Range + Hum Size + Lat. + Forag. + B mass + Hab. + Elev. + Mig. | 0.105 | 0.002 | 0.091 | 0.081 | 0.048 | 0.421 | 0.178 | 0.105 |
| Range + B mass + Elev. + Mig. + Lat. + Forag. + Hab. + Hum Size | 0.105 | 0.02 | 0.017 | 0.167 | 0.029 | 0.069 | 0.481 | 0.005 |
| Mig. + Forag. + B mass + Lat. + Hab. + Range + Elev. + Hum Size | 0.13 | 0.084 | 0.021 | 0.022 | 0.497 | 0.422 | 0.029 | 0.005 |
| Mig. + B mass + Hum Size + Forag. + Lat. + Hab. + Elev. + Range | 0.13 | 0.021 | 0.005 | 0.075 | 0.178 | 0.556 | 0.146 | 0.491 |
| Mig. + B mass + Lat. + Forag. + Hum Size + Range + Elev. + Hab. | 0.13 | 0.021 | 0.025 | 0.069 | 0.001 | 0.545 | 0.132 | 0.584 |
| Mig. + Elev. + Forag. + B mass + Hum Size + Lat. + Hab. + Range | 0.13 | 0.017 | 0.078 | 0.017 | 0.006 | 0.206 | 0.62 | 0.491 |
| Hab. + Forag. + Lat. + Elev. + Hum Size + Range + Mig. + B mass | 0.471 | 0.076 | 0.034 | 0.057 | 0.002 | 0.375 | 0.081 | 0.119 |
| Hab. + Elev. + Forag. + Hum Size + Range + Mig. + Lat. + B mass | 0.471 | 0.026 | 0.066 | 0.005 | 0.185 | 0.091 | 0.128 | 0.119 |
| Hab. + Elev. + Mig. + Forag. + Range + Hum Size + Lat. + B mass | 0.471 | 0.026 | 0.1 | 0.063 | 0.214 | 0.009 | 0.128 | 0.119 |
| Hab. + Range + B mass + Mig. + Hum Size + Forag. + Lat. + Elev. | 0.471 | 0.094 | 0.02 | 0.184 | 0.007 | 0.064 | 0.144 | 0.156 |

**Table S7. Sensitivity analyses show that strong predictors are robust to phylogenetic signal (λ), whereas weak predictors are not.** Sensitivity analyses of the male PGLS model evaluated under fixed values of Pagel’s λ ranging from 0 to 1. For each predictor, partial $R^{2}$, F- and Z-statistics, and p-values are reported. Humerus size and foraging strategy retain consistent effect sizes and statistical support across λ values, whereas predictors with weaker effects show greater variability in statistical support.

| Predictor | R^2^ | F | Z | p-value | Lambda | Predictor | R^2^ | F | Z | p-value | Lambda |
| --- | --- | --- | --- | --- | --- | --- | --- | --- | --- | --- | --- |
| Humerus size | 0.038 | 3.28 | 3.053 | 0.004 | 0 | Habitat | 0.025 | 0.996 | 0.066 | 0.471 | 0.5 |
| Body mass | 0.022 | 1.929 | 1.96 | 0.027 | 0 | Range | 0.016 | 1.261 | 0.76 | 0.227 | 0.5 |
| Foraging | 0.04 | 1.736 | 2.134 | 0.019 | 0 | Latitude | 0.009 | 0.687 | -0.826 | 0.791 | 0.5 |
| Migration | 0.034 | 0.972 | -0.018 | 0.501 | 0 | Elevation | 0.012 | 0.964 | 0.094 | 0.456 | 0.5 |
| Habitat | 0.025 | 1.069 | 0.325 | 0.381 | 0 | Humerus size | 0.033 | 2.662 | 2.733 | 0.003 | 0.75 |
| Range | 0.012 | 1.007 | 0.147 | 0.449 | 0 | Body mass | 0.021 | 1.689 | 1.588 | 0.056 | 0.75 |
| Latitude | 0.007 | 0.618 | -1.049 | 0.853 | 0 | Foraging | 0.036 | 1.465 | 1.594 | 0.058 | 0.75 |
| Elevation | 0.026 | 2.251 | 2.198 | 0.016 | 0 | Migration | 0.041 | 1.09 | 0.497 | 0.32 | 0.75 |
| Humerus size | 0.038 | 3.258 | 3.044 | 0.004 | 0.01 | Habitat | 0.024 | 0.972 | -0.005 | 0.496 | 0.75 |
| Body mass | 0.022 | 1.932 | 1.965 | 0.026 | 0.01 | Range | 0.016 | 1.302 | 0.856 | 0.201 | 0.75 |
| Foraging | 0.039 | 1.691 | 2.048 | 0.024 | 0.01 | Latitude | 0.009 | 0.686 | -0.841 | 0.807 | 0.75 |
| Migration | 0.034 | 0.974 | -0.012 | 0.502 | 0.01 | Elevation | 0.012 | 0.985 | 0.165 | 0.437 | 0.75 |
| Habitat | 0.025 | 1.066 | 0.313 | 0.386 | 0.01 | Humerus size | 0.034 | 2.798 | 2.842 | 0.005 | 0.9 |
| Range | 0.012 | 1.013 | 0.163 | 0.441 | 0.01 | Body mass | 0.019 | 1.58 | 1.38 | 0.077 | 0.9 |
| Latitude | 0.007 | 0.621 | -1.037 | 0.852 | 0.01 | Foraging | 0.04 | 1.642 | 1.969 | 0.027 | 0.9 |
| Elevation | 0.025 | 2.16 | 2.11 | 0.018 | 0.01 | Migration | 0.042 | 1.139 | 0.662 | 0.259 | 0.9 |
| Humerus size | 0.038 | 3.178 | 3.009 | 0.004 | 0.05 | Habitat | 0.023 | 0.952 | -0.044 | 0.523 | 0.9 |
| Body mass | 0.023 | 1.939 | 1.98 | 0.027 | 0.05 | Range | 0.015 | 1.227 | 0.701 | 0.251 | 0.9 |
| Foraging | 0.037 | 1.565 | 1.79 | 0.036 | 0.05 | Latitude | 0.009 | 0.714 | -0.727 | 0.768 | 0.9 |
| Migration | 0.035 | 0.98 | 0.021 | 0.489 | 0.05 | Elevation | 0.014 | 1.113 | 0.513 | 0.309 | 0.9 |
| Habitat | 0.025 | 1.054 | 0.27 | 0.399 | 0.05 | Humerus size | 0.036 | 3.009 | 2.896 | 0.003 | 0.95 |
| Range | 0.012 | 1.038 | 0.229 | 0.417 | 0.05 | Body mass | 0.019 | 1.548 | 1.29 | 0.099 | 0.95 |
| Latitude | 0.007 | 0.633 | -0.996 | 0.84 | 0.05 | Foraging | 0.042 | 1.734 | 2.104 | 0.022 | 0.95 |
| Elevation | 0.022 | 1.864 | 1.783 | 0.042 | 0.05 | Migration | 0.042 | 1.176 | 0.776 | 0.222 | 0.95 |
| Humerus size | 0.037 | 3.087 | 2.968 | 0.005 | 0.1 | Habitat | 0.023 | 0.949 | -0.018 | 0.511 | 0.95 |
| Body mass | 0.023 | 1.941 | 1.988 | 0.027 | 0.1 | Range | 0.014 | 1.146 | 0.519 | 0.31 | 0.95 |
| Foraging | 0.035 | 1.474 | 1.586 | 0.062 | 0.1 | Latitude | 0.009 | 0.773 | -0.493 | 0.679 | 0.95 |
| Migration | 0.036 | 0.991 | 0.068 | 0.471 | 0.1 | Elevation | 0.014 | 1.199 | 0.706 | 0.236 | 0.95 |
| Habitat | 0.025 | 1.042 | 0.226 | 0.418 | 0.1 | Humerus size | 0.056 | 4.896 | 3.003 | 0.011 | 0.99 |
| Range | 0.013 | 1.071 | 0.31 | 0.382 | 0.1 | Body mass | 0.018 | 1.541 | 1.2 | 0.113 | 0.99 |
| Latitude | 0.008 | 0.645 | -0.956 | 0.833 | 0.1 | Foraging | 0.04 | 1.732 | 1.897 | 0.034 | 0.99 |
| Elevation | 0.019 | 1.602 | 1.429 | 0.086 | 0.1 | Migration | 0.044 | 1.271 | 0.993 | 0.159 | 0.99 |
| Humerus size | 0.035 | 2.869 | 2.849 | 0.006 | 0.25 | Habitat | 0.021 | 0.92 | -0.042 | 0.513 | 0.99 |
| Body mass | 0.024 | 1.923 | 1.964 | 0.026 | 0.25 | Range | 0.012 | 1.006 | 0.198 | 0.423 | 0.99 |
| Foraging | 0.034 | 1.366 | 1.317 | 0.097 | 0.25 | Latitude | 0.012 | 1.007 | 0.235 | 0.411 | 0.99 |
| Migration | 0.038 | 1.02 | 0.204 | 0.434 | 0.25 | Elevation | 0.015 | 1.322 | 0.91 | 0.17 | 0.99 |
| Habitat | 0.025 | 1.019 | 0.141 | 0.438 | 0.25 | Humerus size | 0.098 | 9.36 | 3.129 | 0.005 | 1 |
| Range | 0.014 | 1.157 | 0.518 | 0.301 | 0.25 | Body mass | 0.017 | 1.575 | 1.205 | 0.119 | 1 |
| Latitude | 0.008 | 0.669 | -0.877 | 0.81 | 0.25 | Foraging | 0.034 | 1.641 | 1.595 | 0.06 | 1 |
| Elevation | 0.015 | 1.183 | 0.659 | 0.247 | 0.25 | Migration | 0.045 | 1.419 | 1.26 | 0.105 | 1 |
| Humerus size | 0.033 | 2.671 | 2.725 | 0.005 | 0.5 | Habitat | 0.018 | 0.866 | -0.184 | 0.584 | 1 |
| Body mass | 0.023 | 1.835 | 1.824 | 0.036 | 0.5 | Range | 0.01 | 0.929 | 0.039 | 0.491 | 1 |
| Foraging | 0.034 | 1.356 | 1.301 | 0.096 | 0.5 | Latitude | 0.014 | 1.33 | 0.866 | 0.19 | 1 |
| Migration | 0.039 | 1.057 | 0.368 | 0.367 | 0.5 | Elevation | 0.015 | 1.408 | 1.003 | 0.156 | 1 |

**Table S8. Sensitivity analyses reveal interactions between foraging strategy and size-related predictors in males.** Results of interaction sensitivity PGLS models for male humerus shape testing whether scaling varies across foraging strategies. For each model, partial R^2, F- and Z-statistics, and p-values are reported for main effects and interaction terms. Interactions involving foraging strategy show consistent statistical support, whereas other predictors remain weak or non-significant.

| Model | Predictor | R^2^ | F | Z | p-value | Significance |
| --- | --- | --- | --- | --- | --- | --- |
| Body mass * Foraging strategy | Humerus size | 0.087 | 8.96 | 3.547 | 0.002 | ** |
|  | Body mass | 0.012 | 1.185 | 0.606 | 0.277 |  |
|  | Foraging strategy | 0.069 | 3.57 | 2.82 | 0.002 | ** |
|  | Migration | 0.04 | 1.358 | 1.2 | 0.108 |  |
|  | Habitat | 0.02 | 1.017 | 0.259 | 0.387 |  |
|  | Range | 0.01 | 1.061 | 0.317 | 0.374 |  |
|  | Latitude | 0.012 | 1.233 | 0.707 | 0.233 |  |
|  | Elevation | 0.014 | 1.464 | 1.091 | 0.149 |  |
|  | Body mass : Foraging strategy | 0.069 | 3.565 | 2.878 | 0.001 | ** |
| Humerus size * Foraging strategy | Body mass | 0.016 | 1.81 | 1.667 | 0.046 | * |
|  | Humerus size | 0.028 | 3.219 | 2.917 | 0.002 | ** |
|  | Foraging strategy | 0.135 | 7.737 | 4.178 | 0.001 | ** |
|  | Migration | 0.033 | 1.257 | 0.983 | 0.171 |  |
|  | Habitat | 0.016 | 0.937 | -0.064 | 0.531 |  |
|  | Range | 0.009 | 1.044 | 0.283 | 0.398 |  |
|  | Latitude | 0.009 | 1.01 | 0.2 | 0.425 |  |
|  | Elevation | 0.015 | 1.756 | 1.672 | 0.053 | . |
|  | Humerus size : Foraging strategy | 0.134 | 7.734 | 4.185 | 0.001 | ** |

**Table S9. A small number of species exhibit elevated estimates of humerus shape evolutionary rates.** Species identified as having high humerus shape evolutionary rates, with corresponding rate estimates shown. No species were identified as low rate under the applied criteria.

| High-rate outliers | | | |
| --- | --- | --- | --- |
| *Metallura phoebe* | *Elvira chionurus* | *Amazilia yucateca* | *Amazilia lactea* |
| 0.22646258 | 0.03908476 | 0.03491671 | 0.02562061 |
| Low-rate outliers: 0 | | | |

**Sexual dimorphism in humerus shape**

Sexual dimorphism in humerus shape was quantified for 41 species with data available for both sexes. The mean within-species male–female Procrustes distance was 0.086 (SD = 0.023; range = 0.053–0.148), indicating consistent but variable shape differences between sexes across species (see Table S10 below for details). For comparison, typical interspecific Procrustes distances among males were larger. Mean pairwise distances among species within the same foraging strategy were 0.108 (SD = 0.026; 264 pairwise comparisons), while mean pairwise distances among species belonging to different foraging strategies were 0.109 (SD = 0.025; 556 pairwise comparisons). Thus, average sexual dimorphism was smaller than typical among-species variation in male humerus shape, regardless of whether species shared the same foraging strategy.

**Relevance for male-based analyses**

This analysis shows that sexual dimorphism in humeral shape is present but moderate in magnitude. Importantly, the average male–female difference within species does not exceed the typical interspecific variation observed among males. Consequently, sexual dimorphism is unlikely to drive the primary structure of the male humeral morphospace or to account for patterns attributed to ecological or functional differentiation among species. These results support the analytical focus on males in the main text, indicating that male-based comparisons capture biologically meaningful interspecific variation rather than artefacts arising from unaccounted sexual dimorphism.

**Table S10. Sexual dimorphism in hummingbird humerus shape relative to interspecific variation.** Summary of Procrustes distances quantifying sexual dimorphism and interspecific variation in humerus shape. Sexual dimorphism was measured as the within-species Procrustes distance between male and female species mean shapes, calculated after a joint Generalized Procrustes Analysis to place both sexes in a common shape space (41 species). Interspecific distances were calculated between male species mean shapes only, both within the same foraging strategy and between different foraging strategies, using all pairwise species comparisons. Ratios compare the mean magnitude of sexual dimorphism to the mean interspecific distances, providing context for evaluating whether male–female differences approach or exceed typical among-species variation.

| Comparison | N | Mean | SD | Median | IQR | Min | Max |
| --- | --- | --- | --- | --- | --- | --- | --- |
| Male–female difference (within species; Procrustes distance) | 41 | 0.086 | 0.023 | 0.081 | 0.026 | 0.053 | 0.148 |
| Within-foraging interspecific differences (males; pairwise species distances) | 264 | 0.108 | 0.026 | 0.104 | 0.039 | 0.053 | 0.186 |
| Between-foraging interspecific differences (males; pairwise species distances) | 556 | 0.109 | 0.025 | 0.105 | 0.034 | 0.060 | 0.203 |
| Ratio: (Mean male–female) / (Mean within-foraging interspecific; males) | — | 0.80 | — | — | — | — | — |
| Ratio: (Mean male–female) / (Mean between-foraging interspecific; males) | — | 0.79 | — | — | — | — | — |
